# SMAD2: Regulatory junction of TGF-β and antigen signaling in mast cells

**DOI:** 10.1101/2025.07.27.666827

**Authors:** Gina Bronneberg, Steffen K. Meurer, Marlies Kauffmann, Chao-Chung Kuo, Christian Liedtke, Ralf Weiskirchen, Michael Huber

## Abstract

**Objective:** TGF-β/SMAD signaling controls mast cell (MC) development and exerts anti-inflammatory functions, while antigen (Ag)-triggered FcεRI/MAPK activation commands pro-inflammatory processes. SMAD2 is a signaling hub integrating both, TGF-β/ALK5-mediated SMAD2 C-terminal-and ERK-mediated SMAD2 linker-phosphorylation. Here, we analyzed the role of SMAD2 in TGF-β-and Ag-stimulated signaling, and their cross-talk which regulate MC responses.

**Methods:** The CRISPR-Cas method was used to disable SMAD2 expression in the PMC-306 cell line. The transcriptome of the corresponding cells was analyzed by NGS in homeostatic and stimulated conditions. Activation of signaling intermediates, target gene responses and effector function were analyzed by qPCR, Western blot and ELISAs. The causal association of SMAD2 with the found effects was proven by re-introduction of a tagged SMAD2 in *knock out* cells.

**Results:** The absence of SMAD2 led to increased proliferation and survival, and decreased transcription of target genes like *Smad7* and *Jun* in steady state and upon TGF-β treatment. Intriguingly, SMAD2 was found to regulate TGF-β-mediated SMAD1/5 activation, resulting in augmented expression of *Id3* in SMAD2-deficient MCs. Unexpectedly, we found that SMAD2 is indispensable for Ag-triggered production of pro-inflammatory cytokines, such as IL-6 and TNF. Re-introducing SMAD2 restores these events with varying sensitivity depending on the receptor system triggered.

**Conclusions:** Our findings highlight a previously unrecognized role of SMAD2 as a decisive hub in TGF-β-ALK5-SMAD1/5 and Ag-FcεRI-MAPK signaling, suggesting that pharmacological modulation of SMAD2 activity could be leveraged as regulator of both suppressive and inflammatory MC functions.

## Introduction

Mast cells (MCs) are granulated, hematopoietic cells that develop in characteristic sequential waves from the yolk sac, the fetal liver, and the bone marrow (1). After birth, immature MCs leave the bone marrow and enter their target tissues, where terminal differentiation is facilitated (1–3). MCs belong to the innate immune system and represent a first line of defense for invading pathogens and are key elements of the anaphylactic reaction (4). Challenging MCs with antigen (Ag) causes a sequential, bipartite response pattern. In a first wave preformed constituents of secretory lysosomes are released-instantaneously-(“degranulation”). The cargo composition of those lysosomes is directed by environmental cues of the destination MCs are attracted to (2). In a second-delayed step-mediators are de-novo synthesized, e.g. TNF and IL-6, and released to orchestrate relevant responses of immune cells, such as their recruitment (3). Crucial for the regulation of those effector functions by the Ag-FcεRI-axis is the activation of MEK/ERK and NFΚB (3,5).

In contrast to Ag, TGF-β mainly impacts MC proliferation and regulates MC phenotypical maturation by inducing a mucosal-type gene signature in order to prepare cells for the Ag-triggered release reaction (6–8). TGF-β binds to heterogeneous transmembrane receptor complexes, composed of type-I and type-II receptors (9,10). Depending on the engaged receptors, i.e. TGF-β-type or BMP-type, downstream signaling is either directed to SMAD2/3 or SMDAD1/5/9 by phosphorylation of their C-terminal domains (11,12). Both SMAD pathways regulate individual sets of target genes, as exemplified by the up-regulation of *Smad7* (common) and *Mcpt1/Chsy1* (mucosal MC-specific), amongst others, by SMAD2/3, and *inhibitors of differentiation* (*Ids)* by SMAD1/5/9 (7,13–16).

In contrast to the linear receptor-mediated signaling (17), SMAD function can be modulated by phosphorylation of their linker domain. Thereby, SMADs are able to integrate TGF-β and MAPK-signaling (18,19). This has been emphasized for the regulation of *Chsy1* in VSMC, where SMAD2 and its ERK1/2 mediated linker-phosphorylation was shown to regulate *Chsy1* expression (20,21). The cross talk of the TGF-β and Ag/FcεRI axis on the level of Ag-mediated target gene expression has recently been shown in human MCs, but the molecular details are yet unknown (8).

In this study, we demonstrate extensive crosstalk of TGF-β and Ag signaling. This is manifested on a molecular level by Ag/ERK1/2-mediated phosphorylation of the SMAD2 linker region, which affects target gene expression exemplified by RT-qPCR and NGS analysis. In addition, using SMAD2-deficient MCs, we proof that TGF-β-induced gene expression functions strictly in a SMAD2-dependent manner, either positively with respect to SMAD2-dependent genes or negatively concerning SMAD1/5-dependent genes. Finally, we provide evidence that SMAD2 promotes IgE-mediated induction of pro-inflammatory cytokine production, adding to the complexity of the functional cross-talk between regulatory TGF-β and inflammatory Ag-signaling.

## Results

### SMAD2 integrates TGF-β and FcεRI/ERK signals via differential phosphorylation of its linker and C-terminus

In murine BMMCs, only TGF-β caused phosphorylation of the SMAD2 C-terminus, whereas Ag stimulation induced strong SMAD2-L phosphorylation (Fig.1A; Suppl. Fig.1A-C). ERK1/2 were markedly phosphorylated upon Ag and SCF, but not upon TGF-β stimulation (Fig.1A; Suppl. Fig.1B). The ALK5 inhibitor SB431542 significantly suppressed TGF-β-induced SMAD2-CT and SMAD2-L phosphorylation (Fig.1B; Suppl. Fig.1D). Ag-and SCF-stimulated P-SMAD2-L was completely inhibited by Trametinib (MEK1/2 inhibitor), whereas TGF-β-induced P-SMAD2-CT/P-SMAD2-L were unaffected (Fig.1C; Suppl. Fig.1E).

**Figure 1.**
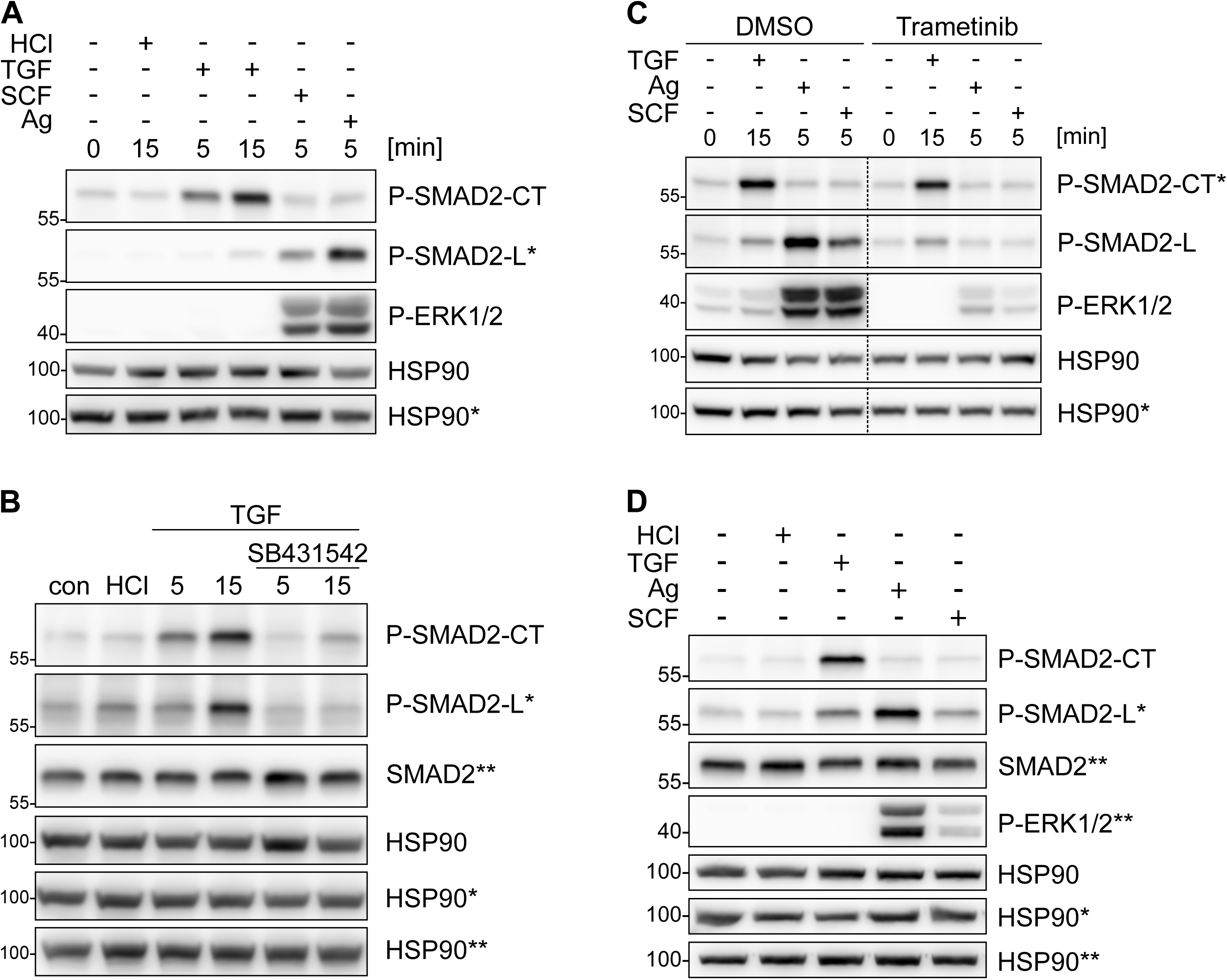
SMAD2-L phosphorylation upon FcεRI crosslinking is dependent on MEK/ERK activation. (A-D) Representative Western blots for P-SMAD2-CT, P-SMAD2-L, SMAD2, P-ERK1/2, and HSP90 as the loading control. Asterisks indicate detection on the same membrane. (A) BMMCs were stimulated for 5 and 15 min with TGF-β, 5 min with SCF, and 5 min with Ag. HCl was used as a control for TGF-β stimulation. n=3 (B) BMMCs were stimulated for 5 and 15 min with TGF-β with pre-incubation of 30 min SB431542 or DMSO control. HCl was used as a control for TGF stimulation. n=3 (C) BMMCs were stimulated for 15 min with TGF-β, 5 min with Ag, and 5 min with SCF with pre-incubation of 30 min Trametinib or DMSO control. Non-relevant lanes were removed from the blot and this is indicated by a dashed line. n=3 (D) PMC-306 cells were stimulated for 15 min with TGF-β, 15 min with Ag, and 15 min with SCF. HCl was used as a control for TGF stimulation. n=3

Observed effects were substantiated in the recently generated, murine peritoneal MC-derived cell line (PMC-306) (22). TGF-β induced significant P-SMAD2-CT, while Ag-stimulated PMC-306 cells showed marked and Trametinib-inhibited P-SMAD2-L (Fig.1D; Suppl. Fig.1F/G). Like in BMMCs, TGF-β did not induce ERK phosphorylation (Fig.1D; Suppl. Fig.1F/G).

In conclusion, Ag stimulation of MCs results in SMAD2-L phosphorylation, which is dependent on the MEK/ERK MAPK pathway. There is no direct cross-activation beside those ligand effector pairs (TGF/SMAD2-CT *vs.* Ag/SCF/MAPK/ SMAD2-L).

### Impact of Ag mediated P-SMAD2-L phosphorylation on TGF-β induced SMAD2 nuclear transfer and *Chsy1* expression

Since SMAD2 is a substrate in Ag signaling, we asked for the functional relevance. Thus, we analyzed if Ag impacts SMAD2 nuclear translocation in absence or presence of TGF-β, and its effect on transcription of TGF-β-responsive genes *Chsy1* (20,23) and *Mcpt2* (24). TGF-β caused a shift of P-SMAD2-CT to the nuclear fraction in BMMCs, which is not affected by Ag (Fig.2A, *left*). Nuclear transfer of Ag-induced P-SMAD2-L only occurred upon co-stimulation with TGF-β (Fig.2A, *right*). TGF-β induced marked transcription of *Chsy1* and *Mcpt2*, which was suppressed by Ag co-stimulation; both mRNAs were not induced by Ag alone (Fig.2B). To determine if Ag-induced MEK/ERK-mediated P-SMAD2-L is involved in this repression, we applied Trametinib, which attenuated the downregulation of *Chsy1* and *Mcpt2* expression in Ag+TGF-β co-stimulated BMMCs without affecting TGF-β stimulation (Fig.2B).

**Figure 2.**
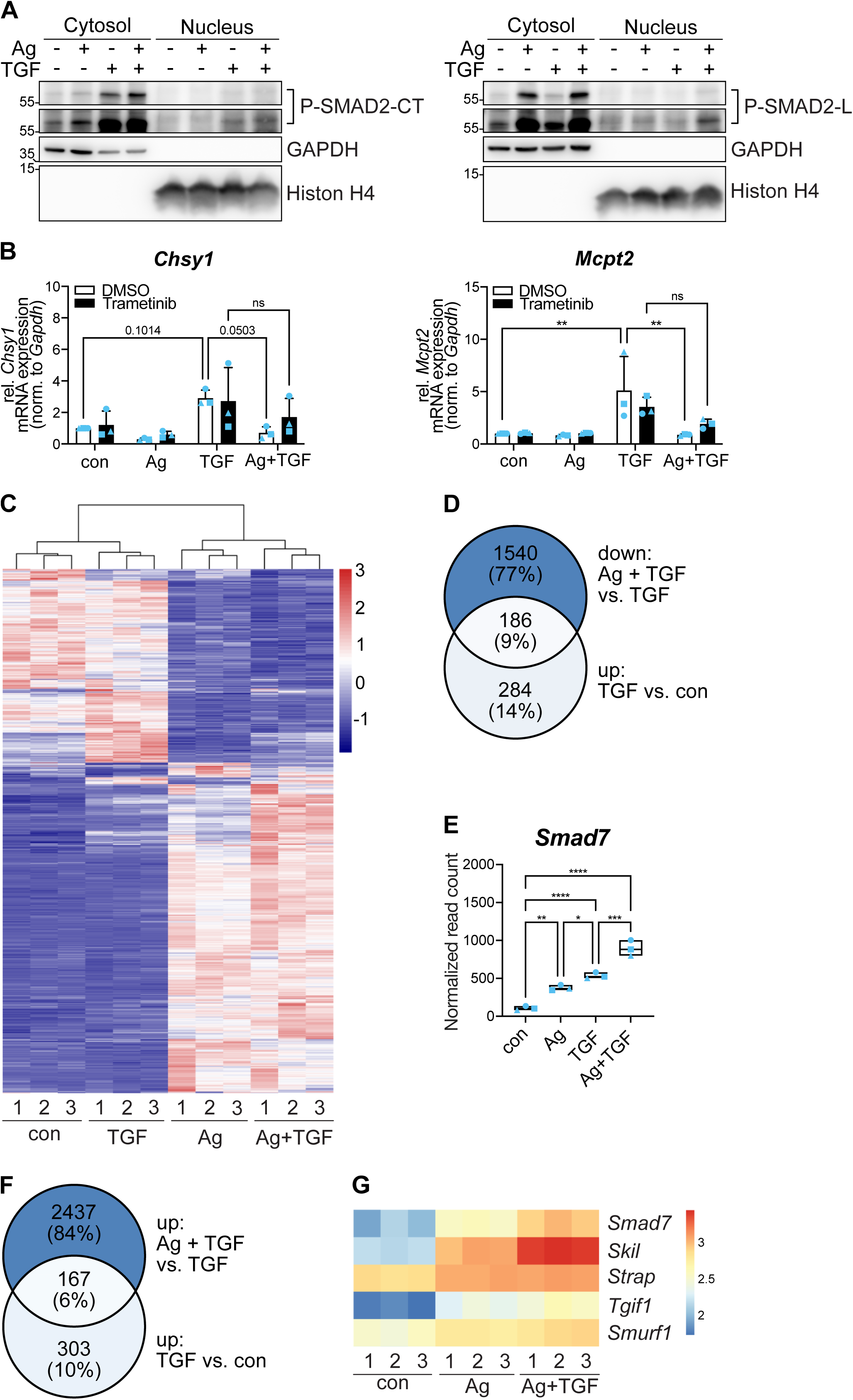
Situation-and gene-dependent mutual regulation of TGF-β and Ag crosstalk. (A) Representative cytosol-nucleus-fractionation for P-SMAD2-CT (*left*) and P-SMAD2-L (*right*) from BMMCs in response to 30 min Ag, TGF-β and Ag+TGF-β. GAPDH served as loading control for the cytosolic fraction and Histon H4 for the nuclear fraction. (B) mRNA expression of *Chsy1* (*left*) and *Mcpt2* (*right*) was assessed by RT-qPCR in BMMCs in response to 90 min Ag, TGF-β and Ag+TGF-β with pre-incubation of 30 min Trametinib or DMSO control. mRNA expression was normalized to *Gapdh*. n=3. (C) Heatmap of genes from BMMCs in response to 90 min TGF-β, Ag and Ag+TGF-β. Rows represent genes and columns represent samples of three independent experiments. The color scale indicates the normalized read counts in log10 scale. (D) Venn diagram illustrating the overlap of genes between two experimental conditions. Each circle represents a set of genes significantly altered. The upper circle represents genes downregulated after Ag+TGF-β stimulation compared to TGF-β treatment, while the lower circle represents genes upregulated after TGF-β stimulation compared to control. The central overlap shows genes regulated across both conditions. (E) Expression of *Smad7* mRNA was identified in BMMCs in response to 90 min Ag, TGF-β, and Ag+TGF-β. The floating bars display the normalized read counts (NGS analysis). n=3 (F) Venn diagram with the upper circle representing genes upregulated after Ag+TGF-β stimulation compared to TGF-β treatment, while the lower circle represents genes upregulated after TGF-β stimulation compared to control. The central overlap shows genes regulated across both conditions. (G) Heatmap of selected negative regulatory genes from BMMCs in response to 90 min con, Ag and Ag+TGF-β. Rows represent genes and columns represent samples of three independent experiments. The color scale indicates the normalized read counts in log10 scale. Bar graphs are shown as mean + SD. *p<0.05, **p<0.01, ***p<0.001, ****p<0.0001 by two-way ANOVA and Tukey’s multiple comparison test (B, C) or one-way ANOVA and Tukey’s multiple comparison test (F).

To shed more light on the impact of Ag on TGF-β-mediated transcription in BMMCs, NGS analysis was performed. A heat map of 1777 differentially expressed genes (DEGs; normalized expression across all samples) allowed clear discrimination between non-and TGF-β-vs. Ag-and Ag+TGF-β-stimulated cells, indicating dominance of Ag-induced over TGF-β-induced signals (Fig.2C). Within these DEGs, 186 genes were found to share the expression pattern of *Chsy1* and *Mcpt2* (Fig.2D). On the other hand, the prototypical TGF-β-target gene Smad7 was increased by Ag signaling itself, which in turn cooperatively promoted TGF-β-mediated expression (Fig.2E; Suppl. Fig.2A). Such pattern was identified for 167 mRNAs, which include further negative regulators of TGF-β-signaling (*Skil*, *Strap*, *Tgif1*, *Smurf1*) (Fig.2F,G; Suppl. Fig.2B).

In conclusion, our data suggest that Ag/MEK/ERK-mediated SMAD2-L phosphorylation can result in transcriptional suppression of several TGF-β-induced genes. Conversely, transcription of a different set of genes, i.e. negative regulators of TGF-β signaling, is enhanced in BMMCs in the presence of Ag. These data imply complex, situation-and gene-specific regulatory mechanisms in TGF-β+Ag-co-stimulated BMMCs.

### Deletion of SMAD2 in mast cells modulates identity, vitality and TGF-β-controlled gene expression

Next, we focused on the role of SMAD2, since SMAD3 is not/hardly expressed in BMMCs and PMC-306 MCs (Suppl. Fig.2C; (16)). Using CRISPR/Cas9, three independent SMAD2-deficient (S2KO) PMC-306 were generated along with their parental cell lines, and confirmed on DNA-, RNA-and protein-level (Suppl. Fig.3A-D). No side effects were expected concerning *Smad1/5* expression due to very low homology of the used gRNA sequence (Suppl. Fig.3E).

**Figure 3.**
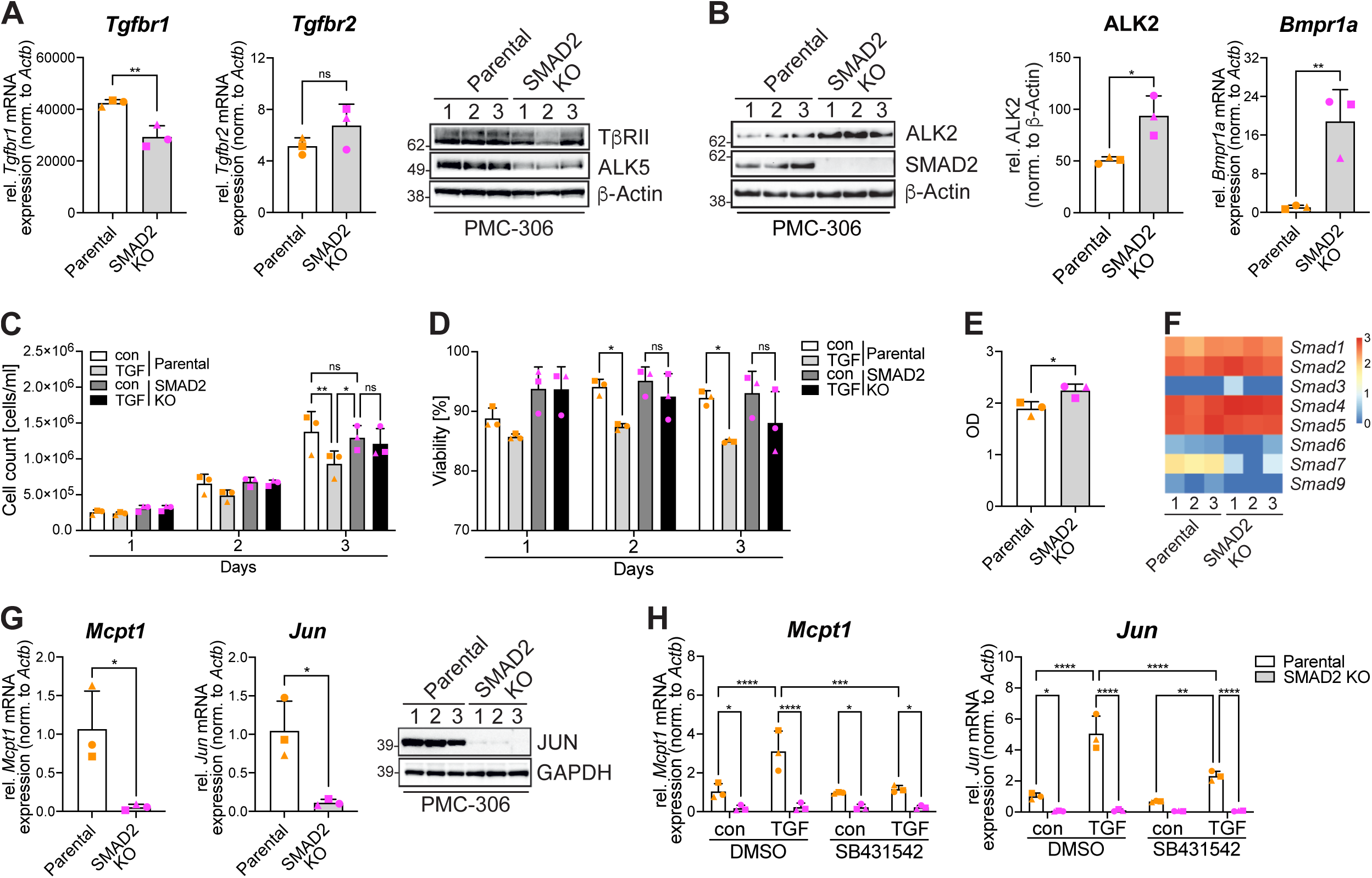
Basic evaluation of SMAD2-deficient (S2KO) MCs. (A) mRNA expression of *Tgfbr1* and *Tgfbr2* was assessed (RT-qPCR) in PMC-306 parental and S2KO cells and normalized to *Actb* (*left panels)*. Expression of ALK5 and TβRII was analyzed by Western Blot, β-Actin served as loading control; n=3 (*right panel*). (B) Expression of ALK2 and SMAD2 was analyzed by immunoblotting, β-Actin served as loading control (*left panel*). Quantification was performed using densitometry; n=3 (*middle panel*). mRNA expression of *Bmpr1a* was assessed (RT-qPCR) in PMC-306 parental and S2KO cells and normalized to *Actb* (*right panel*). (C, D) Cell count (C) and viability (D) of PMC-306 parental and S2KO cells ± TGF-β were measured every 24 h for up to 72 using a CASY cell counter. n=3. (E) The metabolic activity of PMC-306 parental and S2KO cells after 72 h was determined by XTT. n=3. (F) Heatmap of *Smad* genes in PMC-306 parental and S2KO cells. Rows represent genes and columns represent samples. The color scale indicates the normalized read counts in log10 scale. n=3. (G) mRNA expression of *Mcpt1* and *Jun* was assessed (RT-qPCR) in PMC-306 parental and S2KO cells and normalized to *Actb* (*left panels).* Expression of JUN was analyzed by Western blot, GAPDH served as loading control (*right panel*). (H) The mRNA expression of *Jun (left)* and *Mcpt1(right)* was assessed by RT-qPCR in BMMCs in response to 90 min TGF-β with pre-incubation of 60 min SB431542 or DMSO control. The mRNA expression was normalized to *Actb*. n=3. Bar graphs are shown as mean + SD. *p<0.05, **p<0.01, ***p<0.001, ****p<0.0001 by unpaired *t*-test (A, B, E, G) or two-way ANOVA and Tukey’s multiple comparison test (C, D, H).

NGS analysis revealed high expression of TGF-β receptors *Tgfbr1* (encoding ALK5)*, Tgfbr2,* and BMP receptor *Bmpr2* in both parental and S2KO cells (Suppl. Fig.4A). The BMP type-I receptors *Acvr1* (encoding ALK2) and *Bmpr1a* (encoding ALK3) had significantly higher expression levels in S2KO cells (Suppl. Fig.4A). In qPCR and Western blot analyses, *Tgfbr1* showed lower expression at both mRNA and protein levels in S2KO cells (Fig.3A). Expression levels of type-II receptors *Tgfbr2* (Fig.3A), *Bmpr2*, *Acvr2a*, and *Acvr2b* (Suppl. Fig.4B) were similar in both cell types. In line with the NGS data, ALK2 protein and *Bmpr1a* mRNA expression was significantly stronger in S2KO MCs (Fig.3B). FACS analysis revealed higher (FcεRIα) and weaker (KIT) surface expression, consistent with mRNA (*Fcer1a* and *Kit*) and protein (KIT) expression levels (Suppl. Fig.4C), supporting and expanding on data in human skin MCs (25).

**Figure 4.**
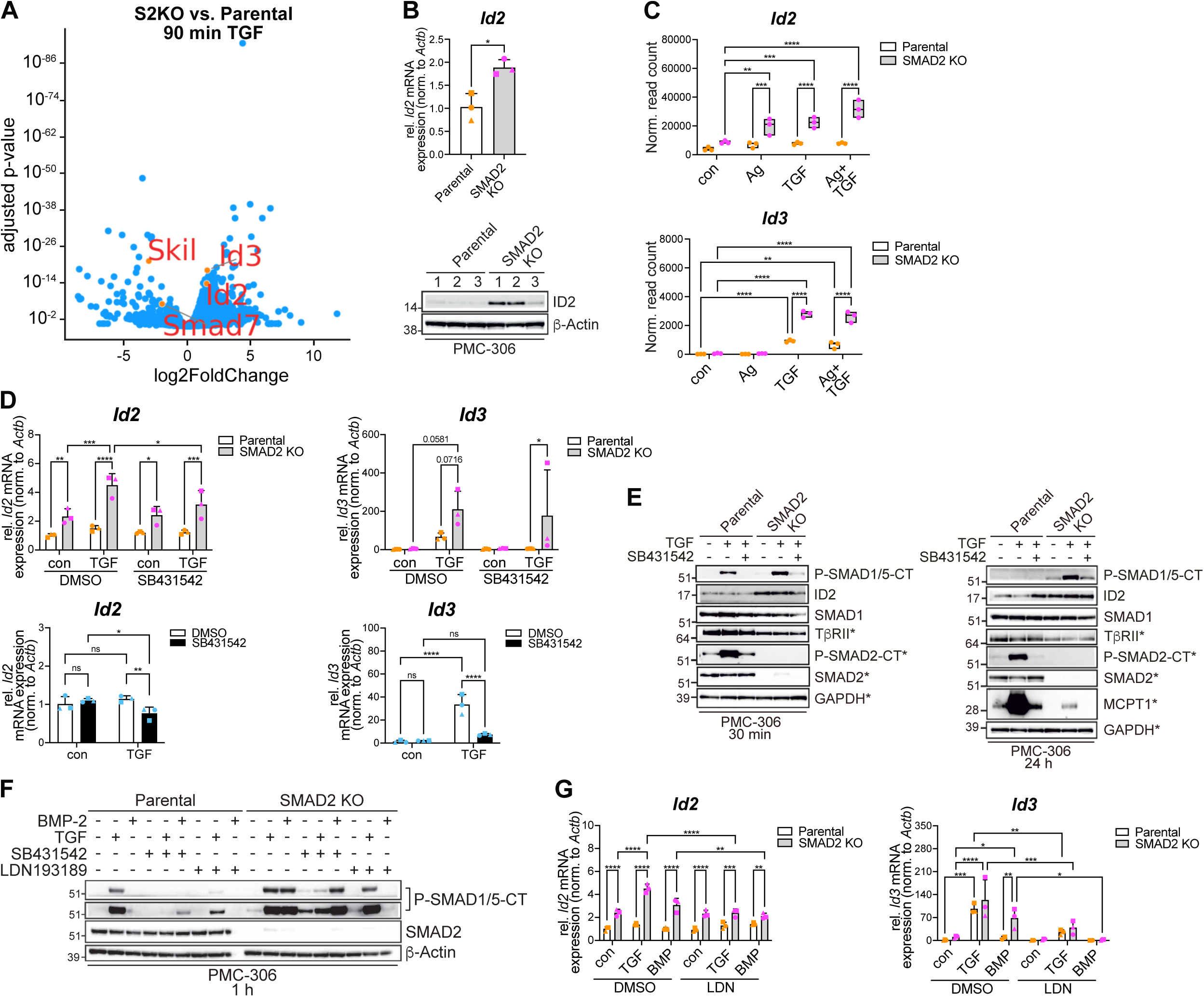
SMAD1/5 activation/function is limited by SMAD2. (A) Volcano plot shows the distribution of genes in PMC-306 S2KO vs. parental cells in response to 90 min TGF-β. The x-axis represents the log2 fold change, y-axis shows the adjusted p-value. Selected genes of interest are indicated (red). (B) mRNA expression of *Id2* was evaluated (RT-qPCR) in PMC-306 parental and S2KO cells and normalized to *Actb (upper)*. Expression of ID2 protein was analyzed by Western Blot. β-Actin served as loading control (*lower*). n=3. (C) Differentially expressed genes from PMC-306 parental and S2KO cells in response to 90 min Ag, TGF-β, and Ag+TGF-β. Floating bars display the normalized read counts of *Id2 (upper)* and *Id3* (*lower*) (NGS analysis). n=3. (D) mRNA expression of *Id2* (*left*) and *Id3 (right)* was assessed (RT-qPCR) in PMC-306 parental and S2KO cells (*upper row*) and BMMCs (*lower row*) in response to 90 min TGF-β with pre-incubation of 60 min SB431542 or DMSO control and normalized to *Actb*. n=3. (E) Representative Western blot for P-SMAD1/5-CT, ID2, SMAD1, TβRII, P-SMAD2-CT, SMAD2, and MCPT1 from PMC-306 parental and S2KO cells in response to 30 min (*left*) or 24 h (*right*) TGF-β with pre-incubation of 60 min SB431542 or DMSO control. Asterisks indicate detection on the same membrane. (F) Representative Western blots for P-SMAD1/5-CT and SMAD2 from PMC-306 parental and S2KO cells in response to 60 min TGF-β or BMP-2 with pre-incubation of 60 min SB431542, LDN193189 (5 μM) or DMSO control. Asterisks indicate detection on the same membrane. β-Actin served as loading control. (G) mRNA expression of *Id2* (*left*) and *Id3* (*right*) was assessed (RT-qPCR) in PMC-306 parental and S2KO cells in response to 60 min TGF-β or BMP-2 with pre-incubation of 60 min LDN193189 (10 μM) or DMSO control and normalized to *Actb*. n=3. Bar graphs are shown as mean + SD. *p<0.05, **p<0.01, ***p<0.001, ****p<0.0001 by unpaired t-test (B), two-way ANOVA and Tukey’s multiple comparison test (C, D (*upper row*), G) or two-way ANOVA and Fisher’s LSD test (D (*lower row*)).

TGF-β stimulation negatively affects proliferation/survival of different cell types including MCs (16,22,26). While parental cells showed reduced cell counts and viability in presence of TGF-β (3 days), this was not observed in S2KO cells (Fig.3C/D). Enhanced proliferation/survival in S2KO cells were accompanied by significant increase in metabolic activity (Fig.3E). We excluded the hypothesis that SMAD3 compensates for SMAD2-deficiency, because both parental and S2KO PMC-306 cells showed only minimal levels of *Smad3* mRNA (Fig.3F; Suppl. Fig.5A) compared to SMAD3-positive primary hepatic stellate cells (Suppl. Fig.5B). In principle, mRNA expression of SMAD family members in parental and S2KO PMC-306 cells was qualitatively and quantitatively comparable to BMMCs, with the exception of *Smad1*/SMAD1 (Suppl. Fig.5C/D), showing reduced expression in S2KO cells.

**Figure 5.**
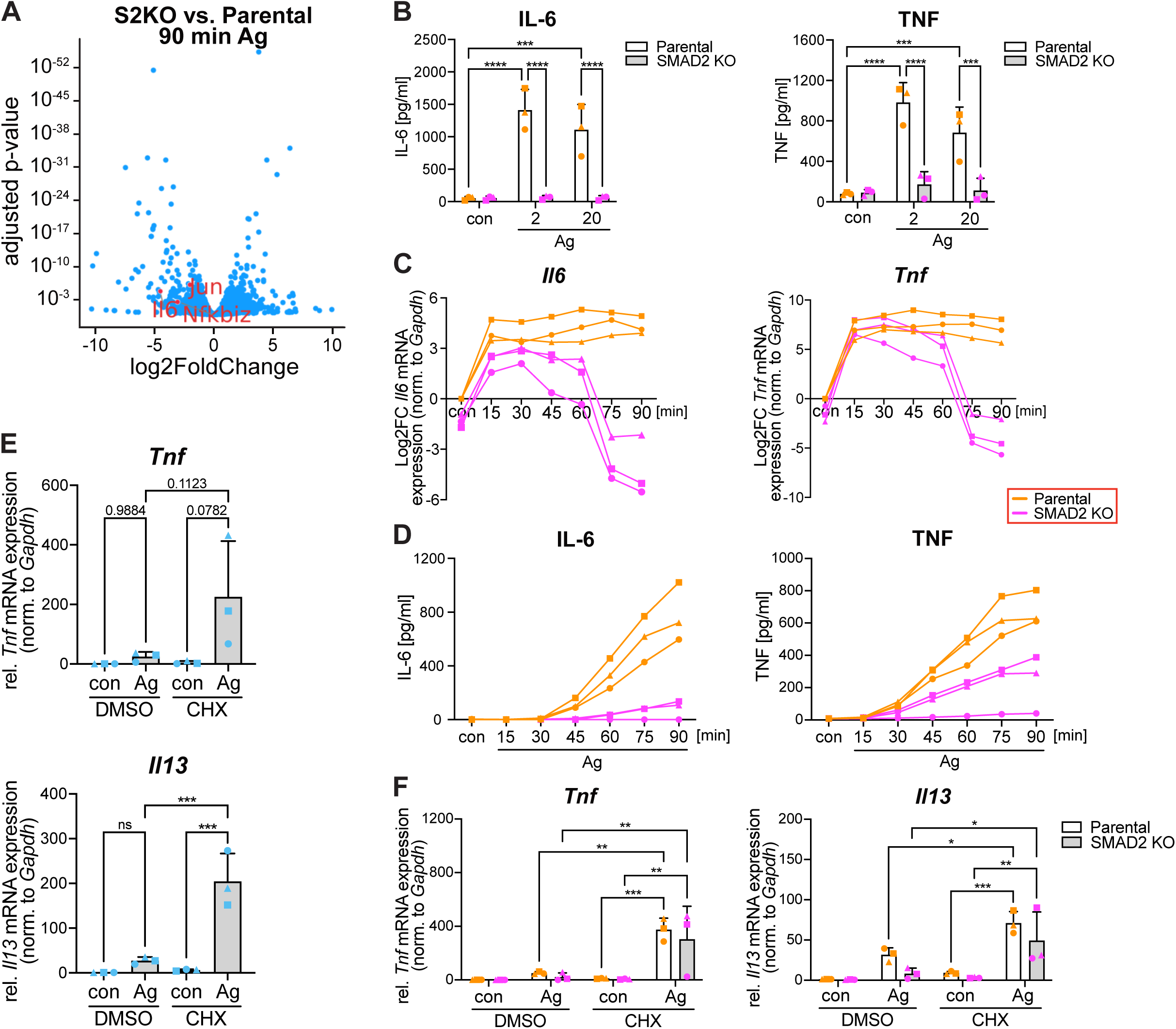
SMAD2 acts as a signaling hub in FcεRI-triggered MCs. (A) The Volcano plot shows the distribution of genes in PMC-306 parental vs. S2KO cells in response to 90 min Ag. The x-axis represents the log2 fold change, the y-axis shows the adjusted p-value. Selected genes of interest are indicated (red). (B) The secretion of IL-6 (*left*) and TNF (*right*) from PMC-306 parental and S2KO cells in response to 3 h Ag (2 and 20 ng/mL) was measured by ELISA. n=3. (C) mRNA expression of *Il6* ((*left*) and *Tnf* (*right*) was assessed (RT-qPCR) in PMC-306 parental and S2KO cells in response to Ag for the indicated time points and normalized to *Gapdh*. Three independent experiments are presented as log2FC. (D) The secretion of IL-6 (*left*) and TNF (*right*) from PMC-306 parental and S2KO cells in response to Ag for the indicated time points was measured by ELISA. n=3. (E) mRNA expression of *Tnf* (*upper*) and *Il13* (*lower*) was assessed (RT-qPCR) in BMMCs in response to 90 min Ag with pre-incubation of 30 min CHX or DMSO control. mRNA expression was normalized to *Gapdh*. n=3. (F) mRNA expression of *Tnf* (*left*) and *Il13* (*right*) was assessed (RT-qPCR) in PMC-306 parental and S2KO cells in response to 90 min Ag with pre-incubation of 30 min CHX or DMSO control. The mRNA expression was normalized to *Gapdh*. n=3. Bar graphs are shown as mean + SD. *p<0.05, **p<0.01, ***p<0.001, ****p<0.0001 by one-way ANOVA and Tukey’s multiple comparison test (E) or two-way ANOVA and Tukey’s multiple comparison test (B, F).

mRNAs of *Mcpt1 and Jun*, and JUN protein were reduced in homeostatic conditions in S2KO MCs (Fig.3G). In line, hardly any *Jun and Mcpt1* mRNA was produced by TGF-β/ALK5 stimulation in S2KO MCs (Fig.3H). In contrast, *Smad7* and *Skil* still appeared slightly inducible by TGF-β/ALK5 in S2KO cells (Suppl. Fig.5E/F).

In conclusion, SMAD2-deficient PMC-306 cells already display perturbations in homeostatic conditions, concerning expression of MC markers, viability and TGF-β target gene expression. Moreover, S2KO MCs showed enhanced proliferation/survival in presence of TGF-β. Furthermore, our data indicated an exclusive SMAD2-dependence of certain TGF-β target genes (e.g. *Jun*), whereas a different set of TGF-β target genes still showed residual responsiveness in the absence of SMAD2 (e.g. *Smad7*).

### SMAD2-deficiency unleashes SMAD1/5 activity and target gene expression

Interestingly, among the genes still responsive to TGF-β we identified DEGs, which are familiar TGF-β/SMAD1/5 targets (e.g. *Id2*, *Id3*) and are even stronger up-regulated in S2KO cells (Fig.4A; Suppl. Fig.6A). These genes were also upregulated by TGF-β in BMMCs (Suppl. Fig.6B). *Id* gene expression is known to be SMAD1/5-dependent (14–16,27), suggesting reinforcement of SMAD1/5 signaling in SMAD2-deficient MCs. Basal expression of *Id2*/ID2 was increased in S2KO vs. parental cells (Fig.4B), and both *Id2* and *Id3* mRNAs were induced stronger by TGF-β in S2KO cells (Fig.4C/D). Increased expression of SMAD1/5-dependent genes in S2KO cells is consistent with enhanced *Acvr1/*ALK2 expression (Fig.3B; Suppl. Fig.4A), as a requirement of two classes of type-I receptors, TGFBR1 (ALK5) and ACVR1 (ALK2), for TGF-β-induced SMAD1/5 phosphorylation has been demonstrated (28). ALK5 involvement in the expression of *Id2* and *Id3* in PMC-306 and BMMC was confirmed (SB431542; ALK5 inhibitor) (Fig.4D).

**Figure 6.**
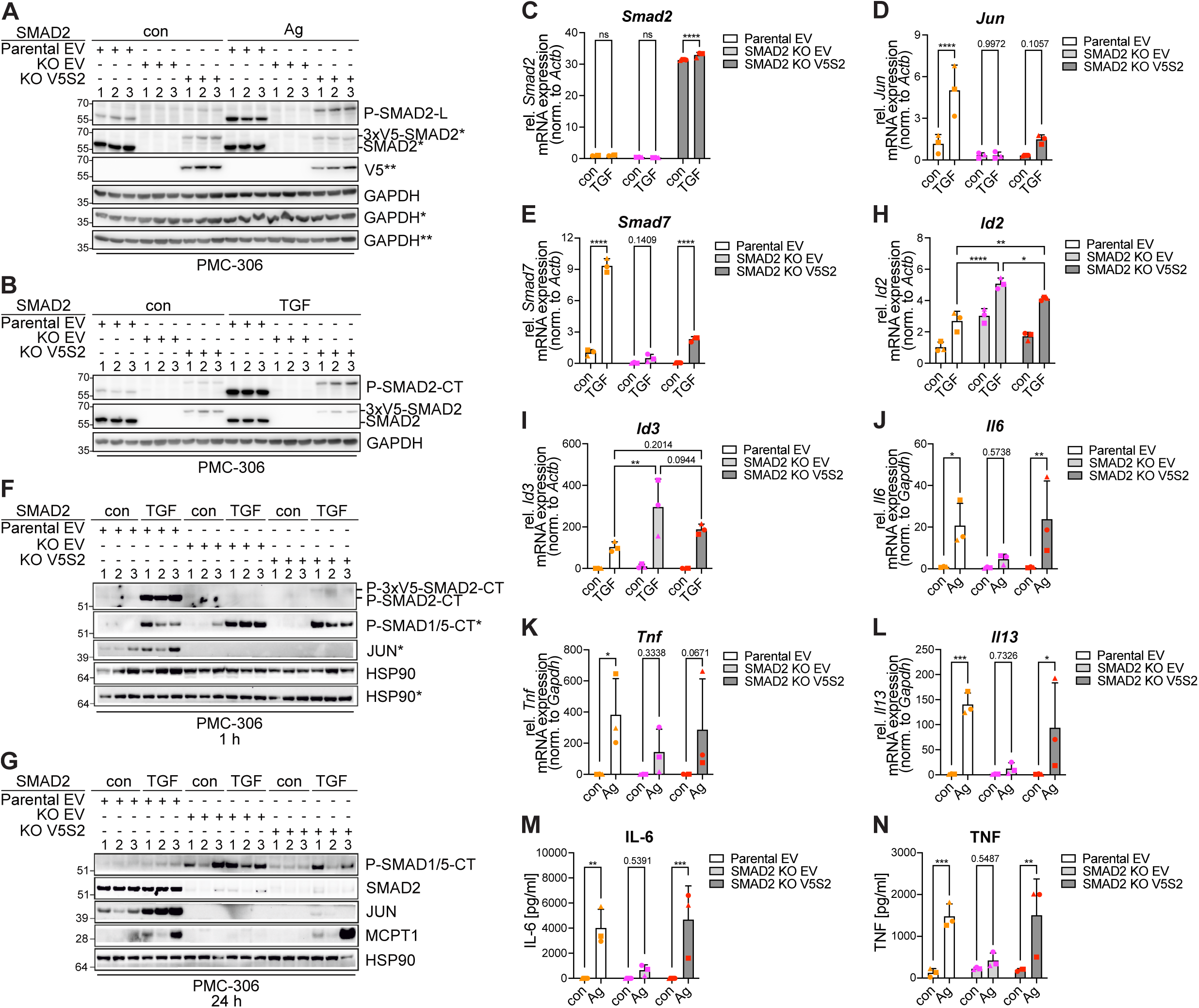
Ag-and TGF-β-induced MC activation exhibit differential sensitivity to SMAD2 expression levels. (A, B, F, G) Western blots for P-SMAD2-CT, P-SMAD2-L, SMAD2, V5, P-SMAD1/5-CT, JUN and MCPT1 from empty vector transfected PMC-306 parental (Parental EV) or S2KO cells (KO EV), and 3xV5-Smad2 transfected S2KO cells (KO V5S2) in response to 15 min Ag (A), 15 min TGF-β (B), 1 h TGF-β (F) and 24 h TGF-β (G). GAPDH and HSP90 served as the loading control. Asterisks indicate detection on the same membrane. (C-E, H-L) mRNA expression of *Smad2* (C), *Jun* (D), *Smad7* (E), *Id2* (h), *Id3* (I), *Il6* (J), *Tnf* (K), and *Il13* (L) was assessed (RT-qPCR) in PMC-306 Parental EV, KO EV and KO V5S2 cells in response to 90 min TGF-β and normalized to *Actb* (C-E, H, I) or *Gapdh* (J-L). n=3. (M, N) The secretion of IL-6 (M) and TNF (N) from PMC-306 Parental EV, KO EV and KO V5S2 cells in response to 3 h Ag was measured by ELISA. n=3. Bar graphs are shown as mean + SD. *p<0.05, **p<0.01, ***p<0.001, ****p<0.0001 by two-way ANOVA and Tukey’s multiple comparison test (C-E, H-N).

Since in S2KO MCs *Id2*/*Id3* expression is increased under basal conditions and/or upon TGF-β stimulation, we asked if this regulation takes place at the transcriptional or the SMAD activation level. SMAD1/5-CT phosphorylation revealed severe kinetic differences between parental and S2KO MCs upon TGF-β stimulation (Fig.4E). P-SMAD1/5-CT intensity appeared already increased in S2KO cells after 30 min, and showed a more digital pattern after 24h, with P-SMAD1/5-CT being not detectable in parental cells. In comparison, P-SMAD2-CT activation remained stable over time in parental cells. Both P-SMAD2-CT and P-SMAD1/5-CT activation were blocked by SB431542 (Fig.4E). This suggests that SMAD2 limits SMAD1/5 activation via a secondary mechanism, i.e. early transcription/translation of (a) repressive factor(s), or downregulation of a BMP-receptor (e.g. ALK2).

To interrogate involvement of type-I BMP receptor(s), we used LDN193189 (ALK2/3 inhibitor) (29). Parental and S2KO PMC-306 cells were stimulated with TGF-β or BMP-2 (strictly SMAD1/5-activating) for 60 min with/without SB431542 or LDN193189 pretreatment, and phosphorylation of SMAD1/5-CT was analyzed. P-SMAD1/5-CT was markedly stronger in TGF-β-vs. BMP-2-stimulated parental cells (Fig.4F). This was not observed in S2KO cells (Fig.4F), suggesting that SMAD2 limits SMAD1/5 activation independent of the stimulus. BMP-2 itself did not induce SMAD2-CT phosphorylation (Suppl. Fig.6C). Both inhibitors considerably attenuated TGF-β-stimulated P-SMAD1/5-CT, matching with data by Ramachandran (28).

qPCR analysis of representative SMAD2-induced (*Jun*) and SMAD2-repressed (*Id2, Id3*) genes confirmed the observed effect of LDN193189 on TGF-β-and BMP-2-induced P-SMAD1/5-CT. LDN193189 did not affect *Jun* (Suppl. Fig. 6D), but significantly decreased *Id2* and even more *Id3* expression in response to TGF-β or BMP-2 (Fig.4G).

In conclusion, SMAD2 inhibits SMAD1/5 activation and corresponding transcription in MCs following TGF-β stimulation, likely through a secondary mechanism. TGF-β stimulation involves a combination of TGFBR1 and possibly ACVR1/ALK2 to regulate activation of the SMAD1/5 pathway.

### SMAD2 constitutes a signaling hub in mast cell activation downstream of the FcεRI

SMAD2 is key in TGF-β-mediated *Smad7* and *Skil* transcription (Suppl. Fig.5E/F). However, Ag stimulation alone also causes weak upregulation of *Smad7* and *Skil*, a response lost in S2KO cells (Suppl. Fig.5E/F). We hypothesized that SMAD2 might also play a positive role in Ag/FcεRI-triggered transcription of target genes other than *Smad7* and *Skil*. Intriguingly, analysis of NGS data from Ag-stimulated parental and S2KO PMC-306 cells revealed further genes with significantly reduced mRNA expression (e.g. *Il6*, *Jun*, and *Nfkbiz*) in S2KO MCs (Fig.5A).

We stimulated parental and S2KO PMC-306 cells with Ag (15/90 min), and analyzed *Il6* and *Tnf* mRNA expression. Whereas there was no difference after 15 min, a significant reduction was observed in S2KO cells after 90 min (Suppl. Fig.7A). Consistently, IL-6 and TNF protein production/secretion was absent in Ag-stimulated S2KO cells (Fig.5B). The signaling defect conveyed by SMAD2-deficiency was increasingly evident starting at approximately 45 min (Fig.5C; Suppl. Fig.7B). Since detectable IL-6 and TNF protein production in parental cells started between 30 and 45 min of Ag stimulation, measurement of these cytokines was difficult in S2KO cells (Fig.5D; Suppl. Fig.7C).

Based on these kinetics, transcription of such genes could depend on SMAD2 regulating immediate-early expression of relevant transcription factors. Therefore, we stimulated BMMCs and parental/S2KO PMC-306 cells with Ag in the presence or absence of cycloheximide (CHX) to inhibit translation of immediate-early proteins, and measured production of *Tnf* and *Il13* mRNA (Fig.5E/F). Indeed, Ag-induced stimulation in the presence of CHX caused dramatic upregulation (“super-induction”) of *Tnf* and *Il13* production, indicating involvement of an immediate-early, suppressively-acting (transcription) factor (Fig.5E/F). In agreement, CHX-pretreatment restored *Tnf* and *Il13* production in S2KO cells (Fig.5F).

In conclusion, IgE-mediated activation of pro-inflammatory genes is supported by SMAD2-dependent homeostatic suppression of (an) attenuating (transcription) factor(s).

### Re-expression of SMAD2 differentially restores mast cell effector functions

To directly connect observed changes with absence of SMAD2, we re-introduced a V5-tagged SMAD2 (V5S2) into S2KO PMC-306 cells. However, re-expression of V5S2 in S2KO cells only resulted in reduced SMAD2 protein levels compared to parental cells, even though mRNA expression was significantly increased (Fig.6A-C; Suppl. Fig.8A). V5S2 appeared to be functional, as TGF-β/Ag stimulation led to expected P-SMAD2-CT/P-SMAD2-L, respectively (Fig.6A/B).

Consistently, TGF-β-mediated expression of *Jun*/*Smad7* revealed only partial reversion by V5S2 (Fig.6D/E). This was confirmed for JUN protein expression (Fig.6F/G). Not all TGF-β-responsive genes were equally (in-) sensitive to weak SMAD2 reconstitution (e.g. MCPT1; Fig. 6G), indicating that protein-type/stability may mask weak reconstitution. Decreased P-SMAD1/5-CT was only observed after prolonged (24 h) TGF-β stimulation (Fig.6G). In line, induction of *Id1*, *Id2* and *Id3* mRNAs was only slightly reduced in TGF-β-stimulated V5S2 vs. S2KO PMC-306 cells (Fig.6H/I; Suppl. Fig.8B).

Interestingly, upon Ag-stimulation, induction of *Il6*, *Il13*, and *Tnf* mRNA as well as IL-6 and TNF protein expression were fully restored (Fig.6J-N). The mechanistic basis of this differential, ligand-dependent effect of incomplete reconstitution is not yet known. However, FcεRI-triggered signaling likely involves more mutually reinforcing elements/pathways that can mask the reduced expression of V5S2, hinting at SMAD2‘s critical role in IgE-dependent proinflammatory processes.

In conclusion, V5S2 was properly phosphorylated upon differential stimulation, leading to a full reversal of diminished FcεRI-mediated cytokine expression. However, TGF-β-mediated target gene expression via SMAD2 or SMAD1/5 was only partially restored. Nonetheless, these experiments support SMAD2’s direct involvement in these regulations, ruling out secondary effects resulting from experimentally-induced long-term SMAD2 deficiency.

## Discussion

In our current study, we investigated the functional interactions during the transition of MCs preparing for (TGF-β) and executing (Ag/FcεRI) effector functions. As a bridging molecule able to receive and integrate signals from both ligands we identified the TGF-β transcription factor SMAD2. Its additional linker-domain phosphorylation by Ag modulates TGF-β-mediated gene responses. To delineate the role of SMAD2 in MCs in detail, we generated a CRISPR S2KO in PMC-306 cells, which prominently underscores its importance in MCs. Most “common” and “differentiation” related TGF-β gene responses were blunted. Conversely, SMAD1/5 signaling shifts from transient to sustained kinetics in the absence of SMAD2. Most interesting, we found that strength and kinetics of FcεRI-induced production of pro-inflammatory mediators is mitigated, a topic that has not been explored so far. In summary, SMAD2 directly influences differentiation and Ag-triggered effector functions in TGF-β-and/or Ag-activated MCs.

Using S2KO MCs, we identified a group of genes that were transcribed in a SMAD2-dependent manner upon TGF-β stimulation, such as *Chsy1, Jun*, *Skil*, and *Smad7*. However, while *Jun* and *Chsy1* were barely transcribed in response to TGF-β in the absence of SMAD2, *Smad7* and *Skil* appeared responsive even in the absence of SMAD2, albeit at a significantly lower level. This suggests an additional regulatory mechanism beside SMAD2. We demonstrated that this pathway also requires ALK5 and may involve SMAD1/5, which is activated by TGF-β in MCs (16). Interestingly, different SMAD proteins were found to recognize genome-wide 5-bp GC motifs (30), potentially enabling the transcription induction of the same genes by both SMAD2 and SMAD1. Consistent with this, *Smad7* can be transcribed in a SMAD1-and GATA-dependent manner (31). SMAD1/5 are more commonly known for their role in BMP responses. However, their activation by the TGF-β/ALK1 axis in endothelial cells has been demonstrated (32,33). Although ALK1 (ACVRL1) was not detectable in the MCs studied, several studies have shown that TGF-β/SMAD1/5 signaling may be accomplished by different receptor combinations, including ACVR1 (a.k.a. ALK2) (27,31,34,35). In line, ALK2 expression along with responsiveness to BMP-2 increases in the absence of SMAD2, suggesting a functional availability of ALK2 in MCs. The higher ALK2 expression in S2KO cells may not only result in stronger BMP-2 responses, but also strengthen TGF-β-induced SMAD1-CT phosphorylation (36).

Phosphorylation of SMAD1/5 and corresponding target gene expression is transient in MCs and can be increased/extended by the translation inhibitor CHX which implies a secondary regulatory response (16). We demonstrate here that this “deactivation” of SMAD1/5 is lost in the absence of SMAD2, since SMAD1/5 activation/phosphorylation and expression of the target gene *Id2* is extended. The mechanism involved may include increased availability of ALK2 (see above) or changes in the phospho-relay system (37). Several Ser/Thr-phosphatases are capable of dephosphorylating SMAD1-CT (37–39). Regardless of the functional availability, all these phosphatases were expressed at mRNA level, but no significant differences in transcripts were noted between parental and S2KO MCs. In conclusion, we found that SMAD2 can act as a negative regulator of SMAD1/5. However, since cell type-and/or situation-dependent co-factors may be necessary for TGF-β/SMAD2-regulated SMAD1/5 activity, this conundrum cannot be definitely answered at this time.

Ag stimulation reduced TGF-β transcriptional responses, e.g. expression of *Chsy1*. But although receptor-mediated phosphorylation of the SMAD2 C-terminal MH2 residues is crucial for nuclear transfer and transcriptional activity (40), Ag signaling did not impact this C-terminal phosphorylation. It has been shown that a cluster of three Ser residues in the SMAD2 linker region can be phosphorylated by different MAP kinases (41), but its effects may be functionally opposing, having either negative or positive effects on downstream TGF-β signaling (41,42). In vascular smooth muscle cells, various ligands, such as LPS and TGF-β, can induce *Chsy1* through SMAD2 linker phosphorylation (43,44). We did not observe TGF-β-induced activation of ERK1/2, p38, and JNK in MCs, but we show here for the first time that Ag triggers marked SMAD2 linker phosphorylation, which was dependent on MEK/ERK activity. However, this linker-phosphorylated SMAD2 does not translocate into the nucleus in the absence of TGF-β-mediated C-terminal phosphorylation, arguing against an autonomous linker function (41). However, if linker phosphorylation impacts transcriptional regulation by providing docking sites for other regulatory factors is currently being investigated.

Another unexpected finding from our study is the requirement of SMAD2 for prolonged IgE-mediated cytokine production. The Ag-triggered “super-induction” of *Tnf* in presence of CHX suggests limitation of *Tnf* transcription by a SMAD2-dependent suppressive (transcription) factor. Hence, SMAD2 might be part of a double-negative feedback, bi-stable signaling system allowing switch-like regulation of (pro-) inflammatory MC responses (45).

In conclusion, SMAD2 is the central mediator of TGF-β signaling in MCs. Further, it constitutes a multifunctional signaling hub in TGF-β-and Ag-stimulated MCs. Based on our finding of IgE-mediated, MEK/ERK-dependent SMAD2 linker phosphorylation we decided to generate and analyze SMAD2-deficient MCs. This allowed us to decipher the role of SMAD2 in inter-SMAD pathway regulation and the homeostatic control of Ag-triggered cytokine production. NGS and bioinformatics analyses revealed a mutually repressive signaling constellation with SMAD2 as an important element in both TGF-β-and Ag-dependent MC activation. Development of pharmacological SMAD2 inhibitors might allow targeted strengthening of SMAD1/5 function. Moreover, such treatment should help to diminish IgE-mediated inflammation.

## Methods

A full description of all methods is available in the Data S1.

### Animals

All mice used in this study were either on a C57BL/6 or on a mixed C57BL/6 x 129/Sv background. Experiments were conducted in compliance with German legislation governing animal studies and following the principles of laboratory animal care. The mice were housed at the Institute of Laboratory Animal Science, Medical Faculty of RWTH Aachen University, which holds a license for the husbandry and breeding of laboratory animals from the veterinary office of the Städteregion Aachen (administrative district). The institute adheres to a quality management system that is certified according to DIN EN ISO 9001:2015. All protocols are reviewed by a Governmental Animal Care and Use Committee at the Landesamt für Verbraucherschutz und Ernährung, Recklinghausen (LAVE). Mice were sacrificed by cervical dislocation to isolate cells from femurs in order to obtain bone marrow cells, which were differentiated into BMMCs as previously described (46).

### Statistical analysis

All data shown were generated from at least three independent experiments. Statistical analysis and graphing of data were performed using Prism (v10.4.1, GraphPad, SanDiego, USA). All statistical test procedures were done as described in the respective figure legends. p-values were considered statistically significant according to the style in GraphPad Prism (ns: p>0.05, * p<0.05, ** p<0.01, *** p<0.001, **** p<0.0001). The respective number of independent biological replicates per experiment is indicated in the figure legends.

## Acknowledgements

This work was supported by grants from the Deutsche Forschungsgemeinschaft (DFG) (HU794/14-1 to MH, LI1045/6-1 to CL, WE2554/15-1 to RW and ME3431/2-1 to SKM).

## Author contributions

Conceptualization, S.K.M. and M.H. Methodology, S.K.M., C.L., R.W., and M.H.

Investigation, G.B., S.K.M., M.K., and C.-C.K. Validation, G.B. and S.K.M.

Writing – Original draft, S.K.M. and M.H. Writing – Review and editing, G.B., S.K.M., Funding acquisition, S.K.M., C.L., R.W., and M.H. Supervision, M.H.

All authors read and commented on the final version of the manuscript.

Bronneberg et al., Data S1

## Key resources tables

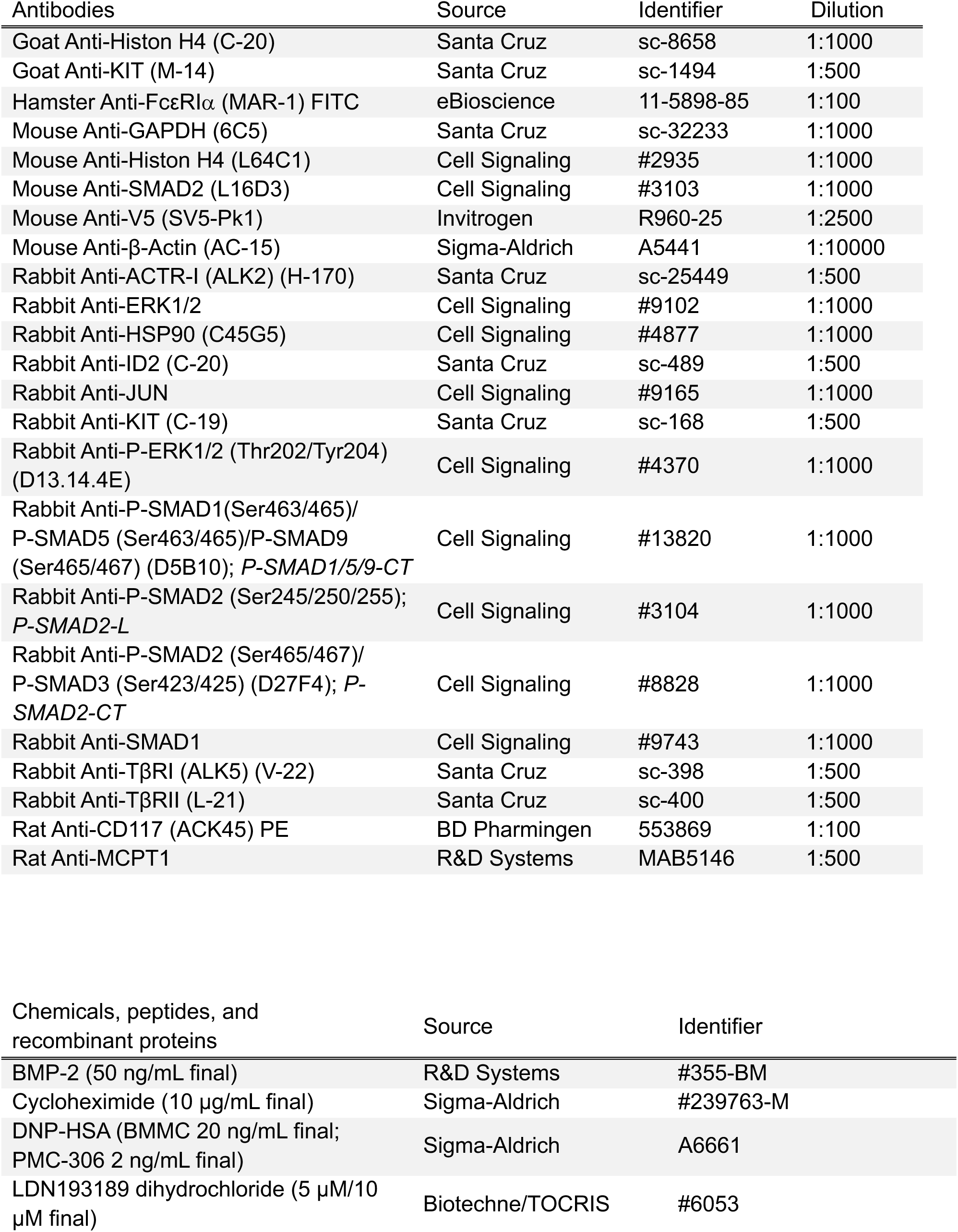
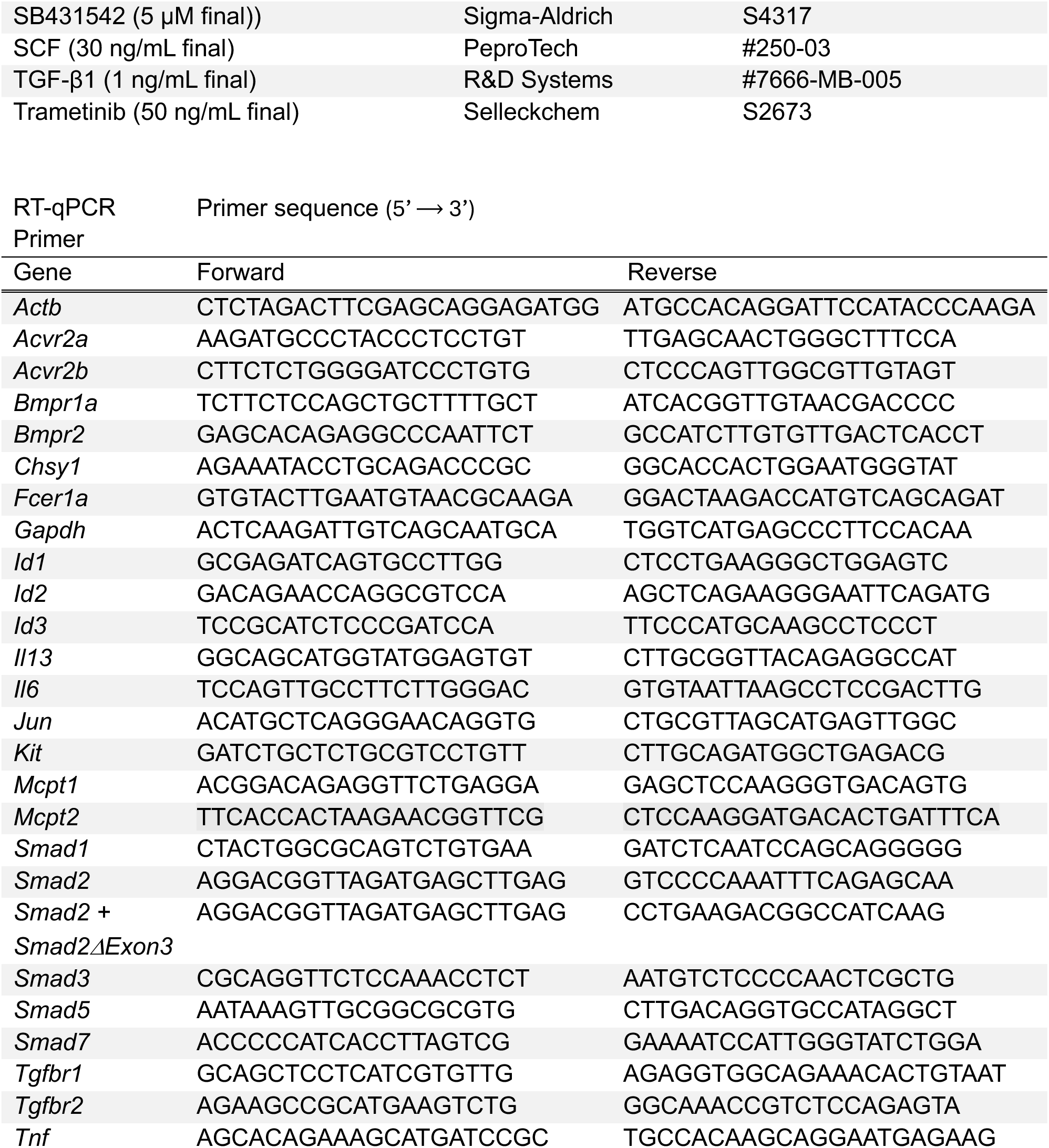

## Experimental model

### Cell culture

All mice used in this study were either on a C57BL/6 background or on a mixed C57BL/6 x 129/Sv background. Experiments were conducted in compliance with German legislation governing animal studies and following the principles of laboratory animal care. The mice were housed at the Institute of Laboratory Animal Science, Medical Faculty of RWTH Aachen University, which holds a license for the husbandry and breeding of laboratory animals from the veterinary office of the Städteregion Aachen (administrative district). The institute adheres to a quality management system that is certified according to DIN EN ISO 9001:2015. All protocols are reviewed by a Governmental Animal Care and Use Committee at the Landesamt für Natur-, Umwelt-und Verbraucherschutz, Recklinghausen (LANUV). No human samples were used, and no experiments were conducted involving living animals. Mice were sacrificed by cervical dislocation to isolate cells from femurs in order to obtain bone marrow cells, which were differentiated into BMMCs as previously described (1).

BMMCs were cultivated as single-cell suspensions in growth medium (RPMI 1640, Gibco Thermo Fisher, #21875-091) containing 15 % heat-inactivated FCS (Capricorn, #FBS-12A or PAN, #P30-3306), 100 units/ml Penicillin and 100 μg/ml Streptomycin (Sigma, #P0781), 10 mM HEPES (Sigma, #H0887), 100 μM β-mercaptoethanol (Sigma, #M6250), and 30 ng/mL IL-3 from X63-Ag8-653 conditioned medium (2). PMC-306 were cultivated in similar growth medium conditions containing 10% heat-inactivated FCS and with the addition of 5 ng/mL SCF derived from cell culture supernatant from CHO cells that were transfected with an expression vector coding for murine SCF (1). The differentiation of BMMCs was evaluated after 4-5 weeks in culture by FACS analysis (see below) and considered successful if more than 95% of BMMCs were positive for MC surface markers FcεRI and KIT. For analysis of BMMCs via Western blot and RT-qPCR (see below), cells were starved (RPMI 1640 + 10% FCS without IL-3) overnight, while BMMCs were not starved for the other experiments. PMC-306 cells were not starved. Additionally, cells were pre-loaded with 0.15 μg/ml IgE (clone SPE-7, Sigma-Aldrich, #D8406) overnight when stimulated with DNP-HSA on the following day.

### CRISPR-Cas9-mediated KO in PMC-306

PMC-306 cells were used to establish a Smad2 KO cell line. A single-guide RNA (sgRNA) complex was created by combining 200 μM crisprRNA against Smad2 (IDT, predesigned Alt-R CRISPR-Cas9 gRNA, sequence: TTCACCACTGGCGGAGTGAA) and 200 μM ATTO550 labeled tracrRNA (IDT, #1075927) at 95 °C for 5 min, followed by cooling to room temperature. Subsequently, 120 pM sgRNA and 104 pM Cas9 enzyme (IDT, #1081058) were mixed in duplex buffer (IDT, #11-01-03-01) and incubated for 15 min at room temperature. 2 x 10^6^ PMC-306 cells along with 1 μl electroporation enhancer (IDT, #1075915) were added and cells were electroporated using the Neon Transfection system (Thermo Fisher, #MPK10096) at 1600 V for 30 mS with one pulse. The cells were then transferred to pre-warmed medium without Penicillin and Streptomycin. The next day, ATTO550 positive cells were individually sorted into 96-well plates by the Flow Cytometry Facility at the Uniklinik Aachen using the BD FACS Aria Fusion cell sorter. Single cells were expanded and subsequently analyzed via Western blot (see below). Smad2 KO clones were sequenced and analyzed with the CRISPR analysis tool by Synthego (3). Three clones with homozygous mutations on both alleles were selected for experiments.

### Retroviral transduction of PMC-306

PMC-306 Smad2 parental and KO cells were used to re-introduce Smad2. Platinum-E cells (Cell Biolabs, #RV-101) were transfected with a construct containing 3xV5-Smad2 and puromycin using the TransIT-LT1 Transfection Reagent (Mirus, #22073360) according to the manufacturer’s instruction and incubated for 24 h. The medium was changed, and the cells were then incubated for an additional 24 h. The retrovirus-containing supernatant was collected, sterile filtered and mixed with the Retro-X Concentrator (Takara, #631455) following the manufacturer’s instructions. The mixture was incubated for at least 6 h at 4 °C and then centrifuged at 1500 x g for 45 min at 4 °C. The pellet was resuspended in PMC-306 growth medium and combined with PMC-306 cells along with 8 μg/ml Polybrene (Merck, #TR-1003-G). On the following day, harvesting of the retrovirus-containing supernatant was repeated, added to the PMC-306 cells and incubated for another 24 h. PMC-306 were centrifuged and resuspended in medium containing 8 μg/ml puromycin (InvivoGen, #ant-pr) for selection. After successful selection, single clones were generated using a dilution series. These clones were propagated and analyzed by Western blot (see below).

### Next generation sequencing

3’mRNA sequencing libraries were prepared using the Lexogen QuantSeq 3’mRNA-Seq v2 Library Prep Kit FWD with Unique Dual Indexes (UDIs) following the manufacturer’s protocol. Prior to library preparation, the concentration of RNA was measured using the Promega Quantus Fluorometer. Additionally, the size distribution of the RNA was assessed using the Agilent TapeStation with an RNA ScreenTape. Quantification and quality assessment were repeated after library preparation, again using the Quantus fluorometer and the Agilent TapeStation with a High Sensitivity D1000 ScreenTape. Libraries were denatured, diluted, and loaded onto a NextSeq High Output v2.5 (75 cycles) flow cell. A 1% PhiX control library was spiked in to improve base calling accuracy. Single-end sequencing was performed with 75 cycles on the Illumina NextSeq platform according to the manufacturer’s instructions.

FASTQ files were generated using bcl2fastq (Illumina). To ensure reproducible analysis, samples were processed using the publicly available nf-core/RNA-seq pipeline version 3.12 (4) implemented in Nextflow 23.10.0 (5) with minimal command. In brief, lane-level reads were trimmed using Trim Galore 0.6.7 (6) and aligned to the mouse genome (GRCm39) using STAR 2.7.9a (7). Gene-level and transcript-level quantification was performed using Salmon v1.10.1 (8). All analysis was conducted using custom scripts in R version 4.3.2 with the DESeq2 v.1.32.0 framework (9).

### ELISA

For the analysis of IL-6 and TNF secretion, MCs were pre-loaded overnight with IgE and ELISA plates (Corning, #9018) were coated with capturing anti-IL-6 (1:250, BD Biosciences, #554400) or anti-TNF (1:250, R&D Systems, #AF-410-NA) antibodies diluted in PBS at 4 °C overnight. Cells were resuspended in stimulation medium (RPMI 1640, #32404 + 0.1 % BSA, Serva, #11930) and the cell number was adjusted to 1.2 x 10^6^ cells/ml. After the cells adapted to 37 °C, they were stimulated for the indicated time points and the supernatant was collected. ELISA plates were washed three times with PBS + 0.1% Tween and blocked with PBS + 2% BSA for 2 h (for IL-6 ELISA) or PBS + 1% BSA + 5 % sucrose for 90 min (for TNF-α ELISA) at room temperature. The plates were washed again and loaded with the cell supernatant (50 μl for IL-6 and 100 μl for TNF-α) as well as a standard dilution for IL-6 (BD Pharmingen, #554582) and TNF (R&D Systems, #410-MT-010). ELISA plates were incubated overnight at 4 °C. Afterwards, the ELISA plates were washed three times and incubated with biotinylated anti-IL-6 (1:500, BD Biosciences, #554402) or anti-TNF (1:250, R&D Systems, #BAF410) antibodies diluted in PBS + 1% BSA for 45 min (for IL-6 ELISA) or 2 h (for TNF-α ELISA) at room temperature. The plates were washed again and incubated with streptavidin alkaline phosphatase (1:1000, BD Pharmingen, #554065) diluted in PBS + 0.5 % BSA for 30 min (for IL-6 ELISA) or 45 min (for TNF-α ELISA) at room temperature. The plates were washed three times and the substrate p-nitro-phenyl-phosphate (Sigma-Aldrich, #S0942, 1 tablet dissolved in 5 ml 2 mM MgCl_2_ in 50 mM sodium carbonate) was added. A plate reader (BioTek Eon) was used to measure OD at 450 nm.

### Flow cytometry

Flow cytometry was used to analyze the surface marker expression of MCs. To confirm the successful differentiation of BMMCs, cells were washed in FACS buffer (PBS + 3 % FCS + 0.1 % sodium azide) and incubated with FITC-coupled anti-FcεRI (1:100, clone MAR-1, eBioscience, #11-5898-85) and PE-coupled anti-CD117 (KIT, 1:100, clone ACK45, BD Pharmingen, #553869) for 30 min at 4 °C in the dark. After another wash, the cells were analyzed using flow cytometry using a FACS Canto II flow cytometer (BD Biosciences). The acquired flow cytometry data were then analyzed using FlowJo software v10.

### Proliferation, Viability and XTT assays

PMC-306 parental and KO cells were seeded at a density of 0.2 x 10^6^ cells/mL and treated as indicated. After 24 hours of incubation in a humidified atmosphere (at 37°C with 5% CO_2_), the cells were resuspended and 50 μl of the cell suspension was diluted in 10 ml of PBS. Cell count and viability were determined using a Casy cell counter from Innovatis. The cell count was monitored over a period of 72 hours.

Metabolic activity was assessed using the XTT cell proliferation kit II (Roche, #11465015001). PMC-306 Smad2 parental and KO cells were plated at a density of 0.3 x 10^6^ cells/ml a 96-well microplate with a final volume of 100 μl. After 72 hours of incubation under humidified culture conditions (37 °C, 5% CO_2_), metabolic activity was measured by adding XTT reagent to each well following the manufacturer’s instructions. Absorbance was then read at 492 nm with a reference wavelength of 650 nm using a plate reader. Total absorbance was calculated by subtracting the absorbance at 650 nm from the absorbance at 492 nm.

### Western blot

Cells were adjusted to a concentration of 1-5 x 10^6^ cells/mL in stimulation medium and stimulated as indicated in the respective experiments. Stimulation was halted by snap-freezing in liquid nitrogen. Pellets were lysed in phosphorylation solubilization buffer containing 50 mM HEPES, 100 mM sodium fluoride, 10 mM sodium pyrophosphate, 2 mM EDTA, 2 mM sodium molybdate, 2 mM sodium orthovanadate, 0.5 % NP-40, 0.3 % sodium deoxycholate, 0.03 % sodium dodecylsulfate, 1 μM PMSF (Sigma, #P7626), 10 μg/mL Aprotinin (Applichem, #A2132), 2 μg/mL Leupeptin (Roth, CN33.1) for 30 min at 4 °C. Lysates were centrifuged for 10 min at 16000 x g and 4 °C, the supernatant was supplemented with loading buffer (30 mM Tris, 2 % glycerol, 2.5 % β-mercaptoethanol, 1% SDS, 0.02% bromophenol blue) and samples were heated at 95 °C for 5 min. Samples were separated on 10 – 15 % SDS polyacrylamide gels using SDS running buffer (25 mM Tris, 192 mM glycine, 0.1 % SDS). Proteins were electroblotted onto PVDF membrane (Roth, #T830.1) and unspecific binding sites were blocked in 5 % non-fat milk powder in PBST (137 mM NaCl, 2.7 mM KCl, 10 mM Na_2_HPO_4_, 2,5 mM KH_2_PO_4_, 0.1 % Tween 20, pH 6.9) for 1h. Most primary antibodies were diluted in 1 % BSA in PBST, while anti-SMAD2 and anti-P-SMAD2 antibodies were diluted in 5% BSA in TBST (20 mM Tris, 150 mM NaCl, 0.05% Tween 20, pH 7.2). Detection was done using horseradish-peroxidase-conjugated secondary antibodies (Dako, #P0448, #P0160, #P0161) and Clarity Max Western ECL Substrate (Biorad, #1705062). Proteins were visualized using a LAS-4000 reader (Fujifilm).

Western blots shown in Figs. 1A-D, 2A, 6A, 6B, and Suppl. Fig. 1B, 1F, 3D, 8A were carried out according to the protocol described above, while Western blots in Figs. 3A, 3B, 4B, 4E, 4F, 6F, 6G, and Suppl. Fig. 4C, 5D, 6C were performed as described previously (10).

### qPCR

Total RNA was isolated from 3-5 x 10^6^ cells using the NucleoSpin RNA Plus Kit (Macherey Nagel, #740984) according to the manufacturer’s instructions. 1 μg of RNA was reverse transcribed using random oligonucleotides (Roche, #11034731.001) and the Omniscript Reverse Transcription Kit (Qiagen, #205113) according to the manufacturer’s instructions. Quantification of transcript expression was carried out using the SensiMix SYBR No-Rox Kit (Bioline, QT650-05) and 10 pmol of specific primers. PCR reactions were conducted on a Rotorgene Q (Qiagen). Transcript expression was normalized to the housekeeping genes *Gapdh* or *Actb* and relative expression was calculated using the delta-C_T_ method (11).

qPCRs shown in Figs. 2B, 2C, 5C, 5E, 5F, 6J-L and Suppl. Figs. 7A, 7B were carried out according to the protocol described above, while qPCRs in Figs. 3A, 3B, 3G, 3H, 4B, 4D, 4G, 6C-E, 6H, 6I, and Suppl. Figs. 2A, 4B, 4C, 5B, 5C, 5E, 6D, 8B were performed as described previously (10).

### Data availability statement

The NGS data sets on TGF-β/Ag-stimulated BMMCs as well as parental and S2KO PMC-306 cells have been deposited in NCBÍs Gene Expression Omnibus and can be found under respective accession codes: GSE????? and GSE?????. These will be made publicly accessible after acceptance of the manuscript.

---

**Bronneberg et al. – Supplement Fig. 1.**
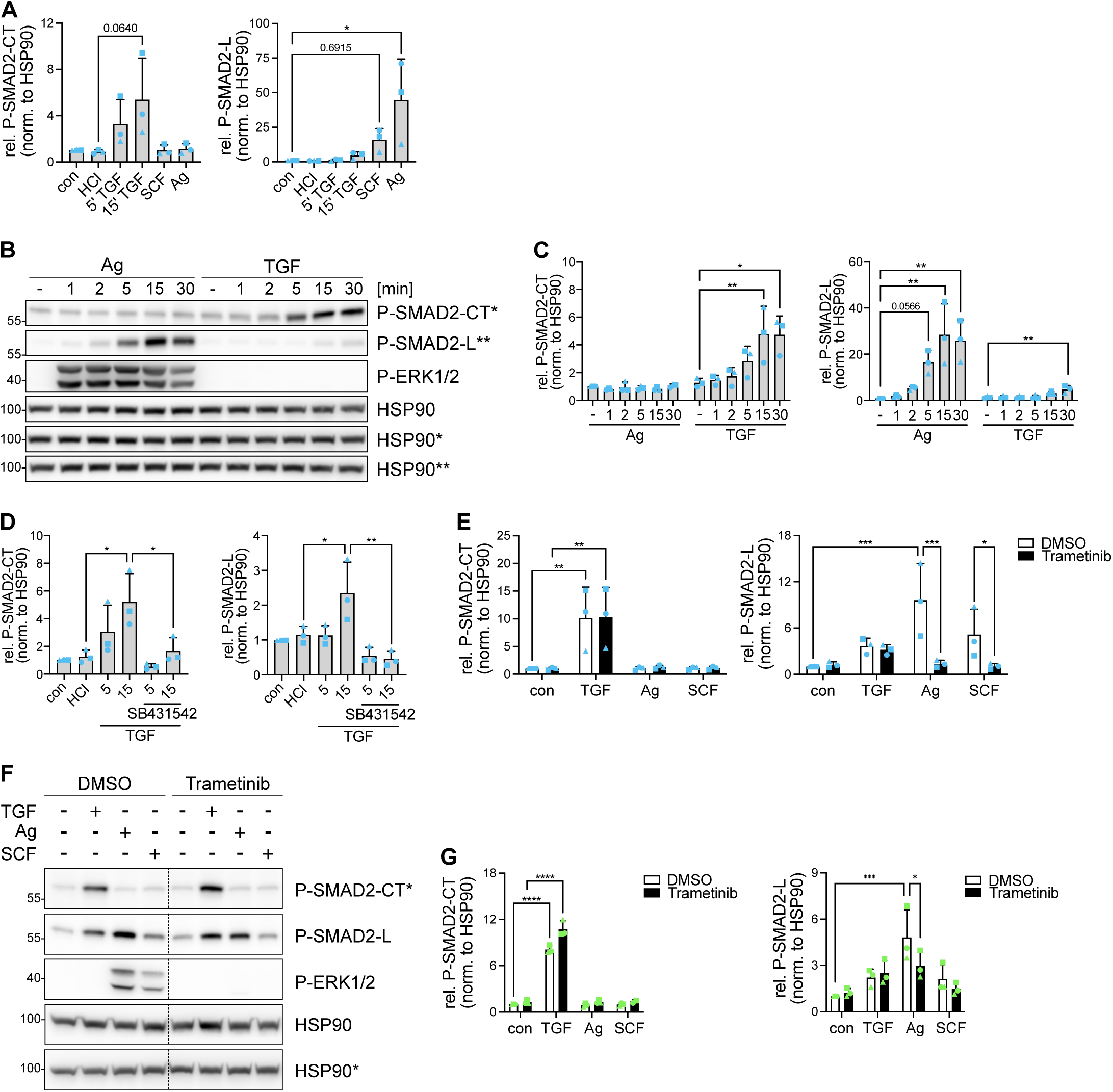
(A) Quantitative analysis of P-SMAD2-CT and P-SMAD2-L protein levels relative to HSP90 (corresponding to Fig. 1A) (B) Representative Western blot for P-SMAD2-CT, P-SMAD2-L, and P-ERK1/2 in BMMCs in response to Ag (20 ng/mL) and TGF-β (1 ng/mL) for the indicated time points. HSP90 served as loading control. Asterisks indicate detection on the same membrane. (C) Quantitative analysis of P-SMAD2-CT and P-SMAD2-L levels relative to HSP90. n=3 (D and E) Quantitative analysis of P-SMAD2-CT and P-SMAD2-L protein levels relative to HSP90. Graphs depicted in (D) correspond to Fig. 1B and graphs depicted in (E) correspond to Fig. 1C. (F) Representative Western blot for P-SMAD2-CT, P-SMAD2-L, and P-ERK1/2 in PMC-306 cells in response to 15 min TGF-β (1 ng/mL), 15 min Ag (2 ng/mL), and 15 min SCF (30 ng/mL) with pre-incubation of 30 min Trametinib (50 nM) or DMSO control. HSP90 served as loading control. Asterisks indicate detection on the same membrane. Non-relevant lanes were removed from the blot indicated by a dashed line. (G) Quantitative analysis of P-SMAD2-CT and P-SMAD2-L levels relative to HSP90. n=3. Bar graphs are shown as mean + SD. *p<0.05, **p<0.01, ***p<0.001, ****p<0.0001 by one-way ANOVA and Tukey’s multiple comparison test (A, C, D), one-way ANOVA and Dunnett’s multiple comparison test (F), or two-way ANOVA and Tukey’s multiple comparison test (B, H).

**Bronneberg et al. – Supplement Fig. 2.**
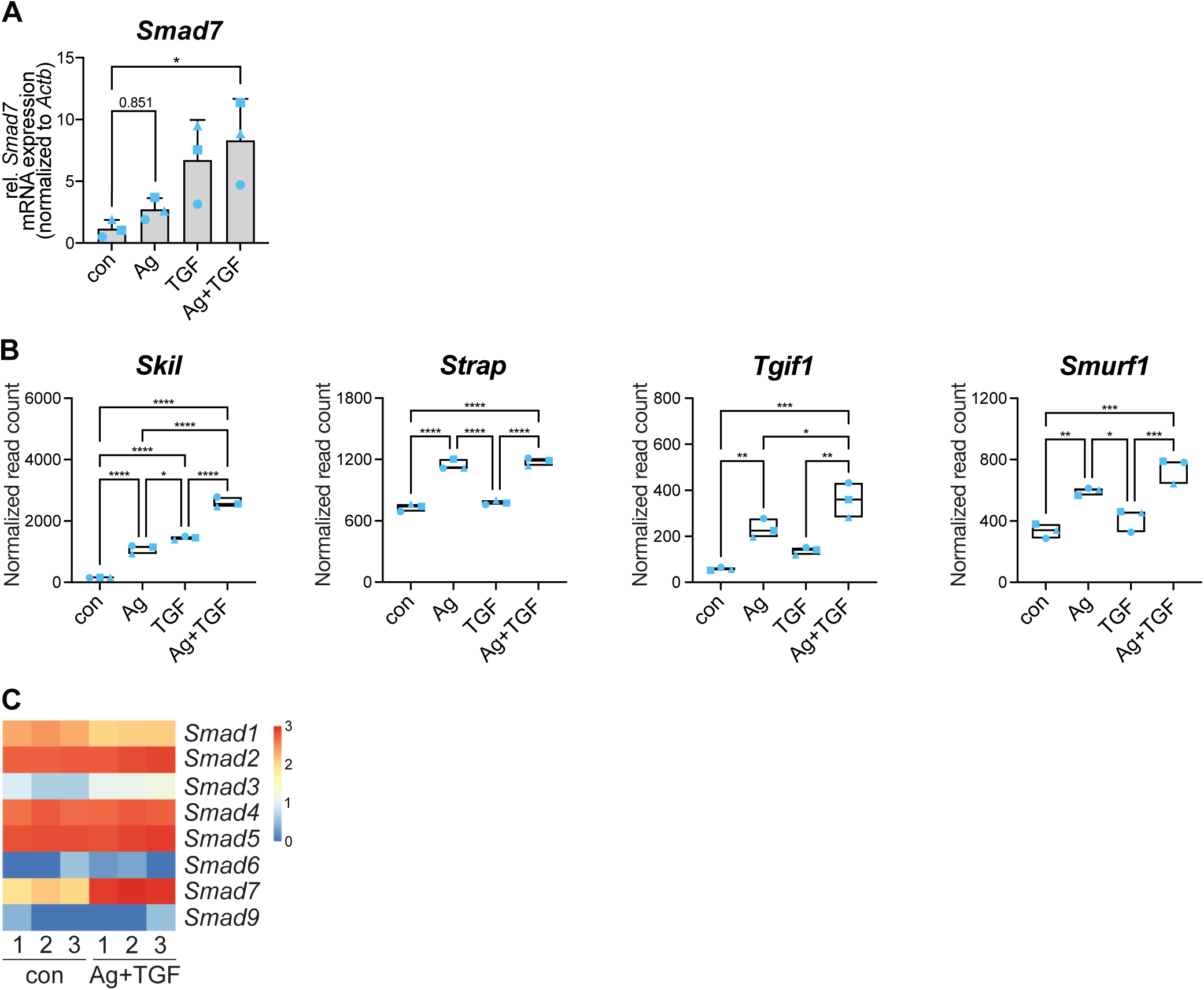
(A) mRNA expression of *Smad7* was assessed by RT-qPCR in BMMCs in response to 90 min Ag, TGF-β and Ag+TGF-β and was normalized to *Actb*. n=3. (B) Differentially expressed genes from BMMCs in response to Ag (2 ng/mL), TGF-β, and Ag+TGF β for 90 min. Floating bars display the normalized read counts (NGS analysis). (C) Heatmap of genes from BMMCs left unstimulated (con) and in response to 90 min Ag (2 ng/mL) + TGF-β. Rows represent genes and columns represent samples of three independent experiments. The color scale indicates the normalized read counts in log10 scale. *p<0.05, **p<0.01, ***p<0.001, ****p<0.0001 by one-way ANOVA and Dunnett’s multiple comparison test (A) or one-way ANOVA and Tukey’s multiple comparison test (B).

**Bronneberg et al. – Supplement Fig. 3.**
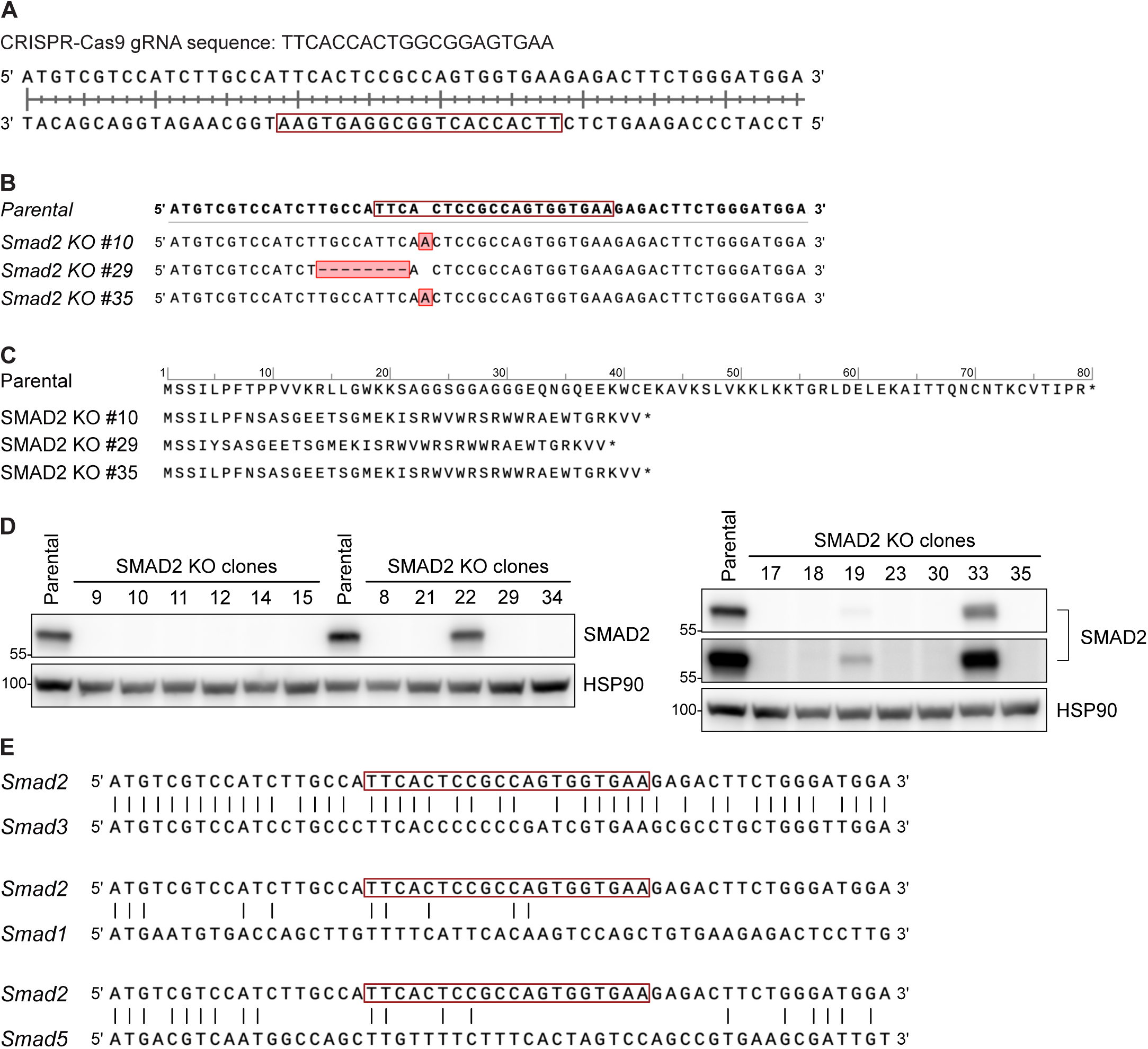
(A) The depicted CRISPR-Cas9 gRNA sequence (red box) of *Smad2* was used to generate PMC-306 SMAD2 KO cells. (B, C) Multiple clones were collected from which number 10, 29 and 35 were sequenced (B) and subsequently selected for experiments. Clone 10 and 35 had the same single nucleotide insertion, while clone 29 had an 8 nucleotide deletion. Those changes result in frame shifts and cause premature stop codons (nonsense mutations) in the amino acid sequence (C, indicated with asterisks). (D) Western blots for SMAD2 using protein extracts from PMC-306 parental and S2KO cells, with HSP90 serving as the loading control. (E) Sequence alignment of *Smad2* with *Smad3*, *Smad1* and *Smad5*, respectively. The red box marks the binding site of the CRISPR-Cas9 gRNA.

**Bronneberg et al. – Supplement Fig. 4.**
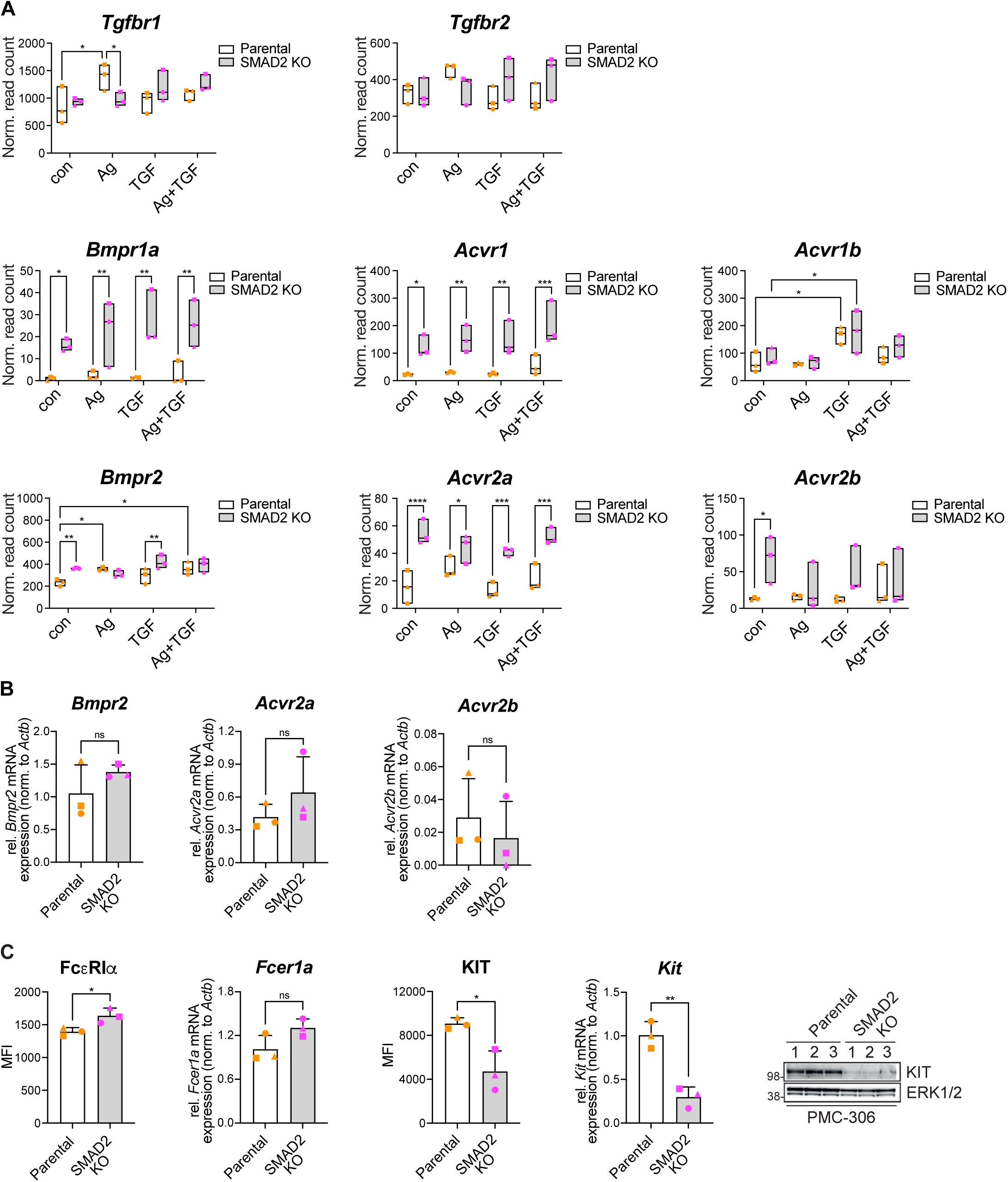
(A) Differentially expressed receptor genes in PMC-306 parental and S2KO cells in response to 90 min of Ag, TGF-β, and Ag + TGF-β. The floating bars display the normalized read counts from NGS analysis, with each replicate represented by a unique symbol. (B) mRNA expression of *Bmpr2, Acvr2a* and *Acvr2b* was assessed (RTqPCR) in PMC-306 parental and S2KO cells and normalized to *Actb*. n=3. (C) Flow cytometric analysis of cell surface expression of FcεR1a and KIT in PMC-306 parental and S2KO cells. Data from three independent experiments are depicted as mean fluorescence intensity (MFI) + SD. mRNA expression of *Fcer1a* and *Kit* was assessed (RT-qPCR) and normalized to *Actb*. Expression of KIT was analyzed by Western blot, ERK1/2 served as loading control. Bar graphs are shown as mean + SD. *p<0.05, **p<0.01, ***p<0.001, ****p<0.0001 by unpaired t-test (B, C) and two-way ANOVA and Tukey’s multiple comparison test (A).

**Bronneberg et al. – Supplement Fig. 5.**
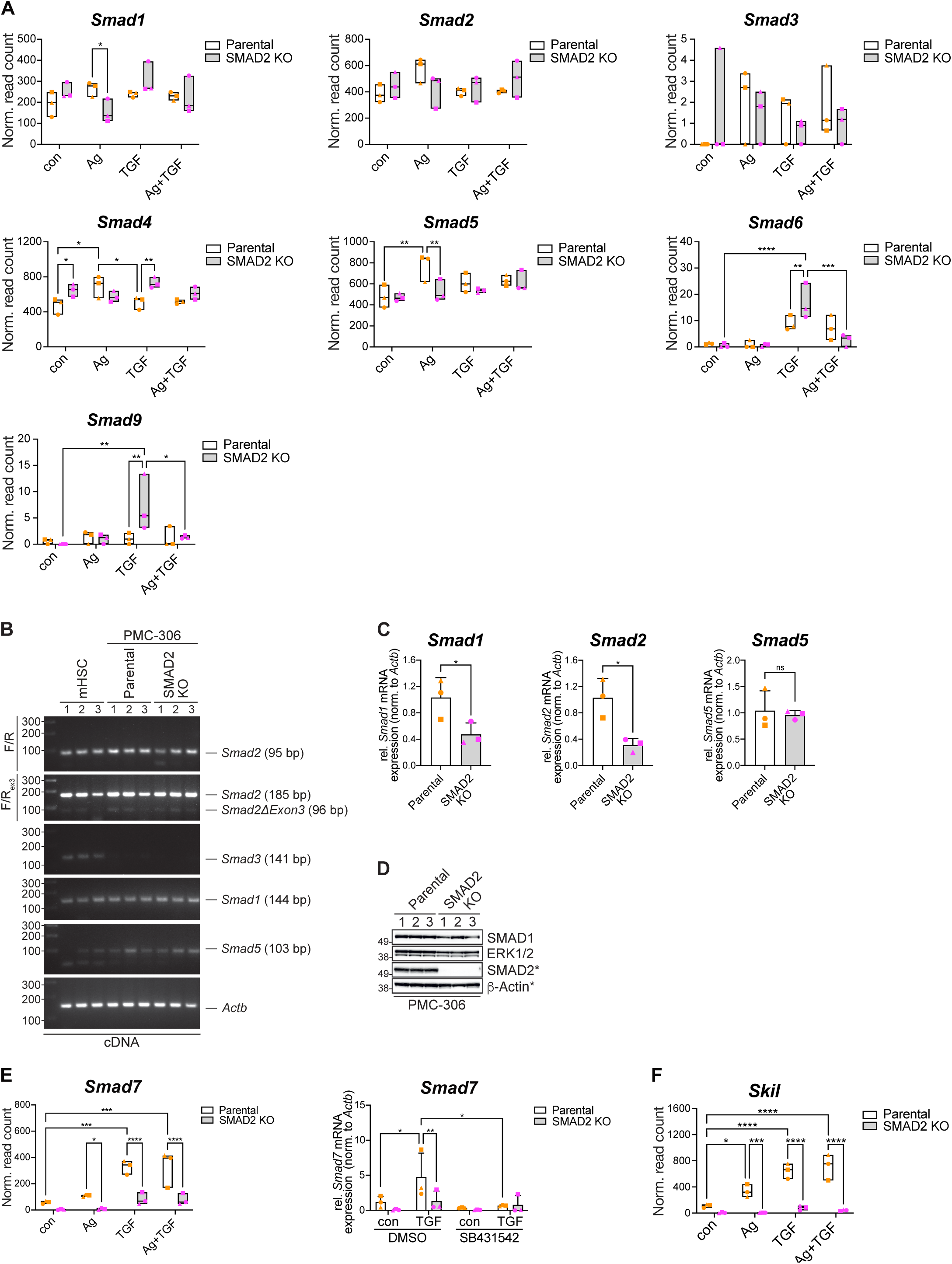
(A) Differentially expressed *Smad* genes in PMC-306 parental and S2KO cells in response to 90 min stimulation with Ag, TGF-β, and Ag + TGF-β. The floating bars display the normalized read counts from NGS analysis. (B) The expression of *Smad2*, *Smad2*D*Exon3*, *Smad3*, *Smad1* and *Smad5* was assessed in murine primary hepatic stellate cells (mHSCs) and PMC-306 parental and S2KO cells using RT-PCR. *Actb* was used as the loading control. (C) mRNA expression of *Smad1, Smad2, and Smad5* was assessed (RT-qPCR) in PMC-306 parental and S2KO cells and normalized to *Actb*. n=3. (D) Western blot analysis for SMAD1 and SMAD2 from PMC-306 parental and S2KO cells. ERK1/2 and β-Actin served as loading controls. Asterisks indicate detection on the same membrane. (E) (Left) Expression of *Smad7* mRNA was identified in PMC-306 parental and S2KO cells in response to 90 min Ag, TGF-β, and Ag+TGF-β. The floating bars display the normalized read counts (NGS analysis). (Right) The mRNA expression of *Smad7* was assessed by RT-qPCR in BMMCs in response to 90 min TGF-β with preincubation of 60 min SB431542 (5 μM) or DMSO control. The mRNA expression was normalized to *Actb*. n=3. (F) Expression of *Skil* mRNA was identified in PMC-306 parental and S2KO cells in response to 90 min Ag, TGF-β, and Ag+TGF-β. The floating bars display the normalized read counts (NGS analysis). Bar graphs are shown as mean + SD. *p<0.05, **p<0.01, ***p<0.001, ****p<0.0001 by unpaired t-test (B) and two-way ANOVA and Tukey’s multiple comparison test (A, E, F).

**Bronneberg et al. – Supplement Fig. 6.**
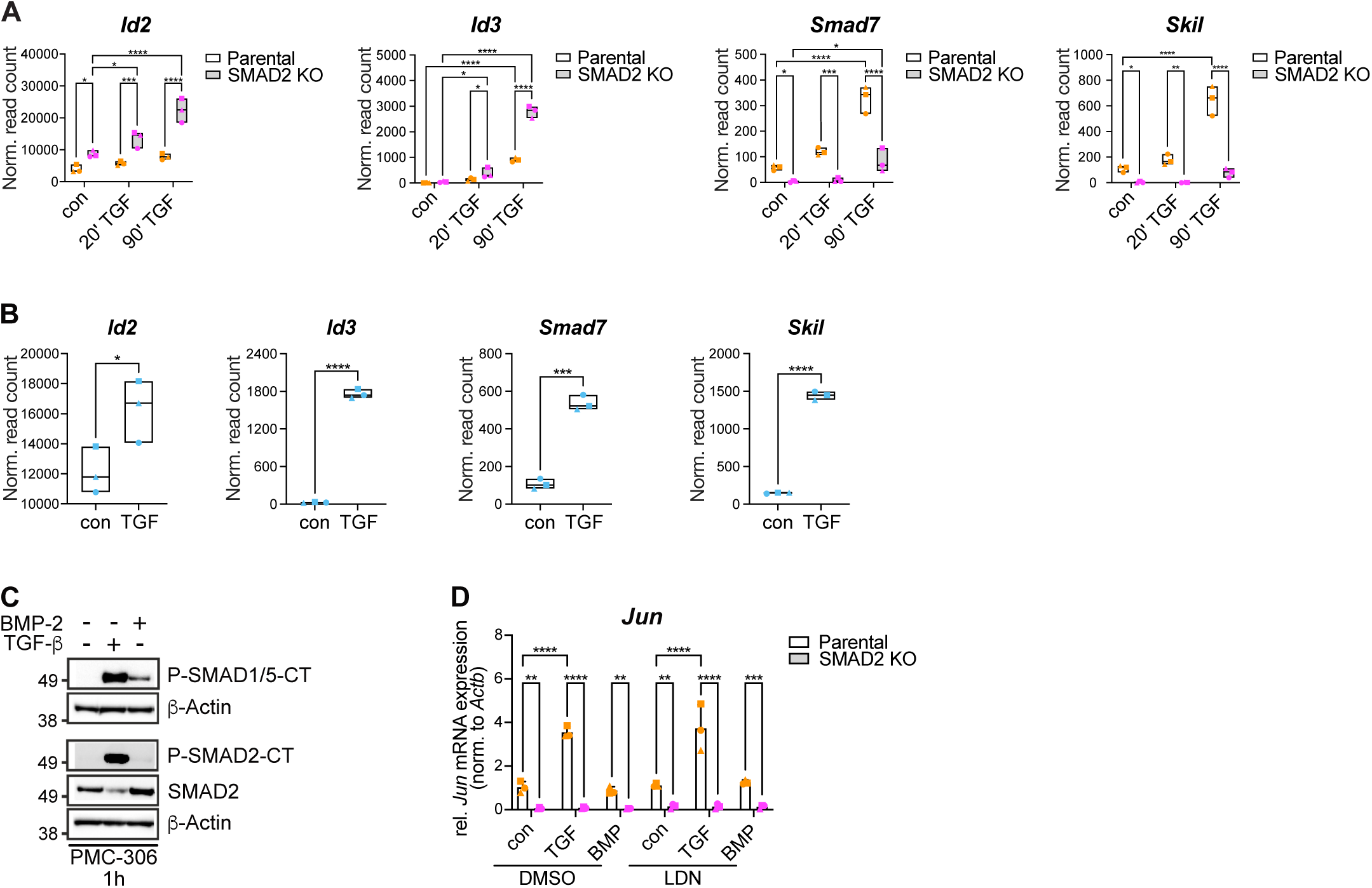
(A) Differentially expressed genes (*Id2*, *Id3*, *Smad7*, *Skil*) from PMC-306 parental and S2KO cells in response to 20 and 90 min TGF-β. The floating bars display the normalized read counts (NGS analysis). (B) Differentially expressed genes (*Id2*, *Id3*, *Smad7*, *Skil*) from BMMCs in response to 90 min TGF-β stimulation. The floating bars display the normalized read counts (NGS analysis). (C) Western blot analysis of PMC-306 cells stimulated or not with TGF-β or BMP-2 (50 ng/mL) for 1 hr and detection of P-SMAD1/5-CT, P-SMAD2-CT and SMAD2, with β-Actin serving as the loading control. (D) mRNA expression of *Jun* was assessed (RT-qPCR) in PMC-306 parental and S2KO cells in response to 60 min TGF-β or BMP-2 with pre-incubation of 60 min LDN193189 (10 μM) or DMSO control and normalized to *Actb*. n=3. *p<0.05, **p<0.01, ***p<0.001, ****p<0.0001 by unpaired t-test (B) and twoway ANOVA and Tukey’s multiple comparison test (A, D).

**Bronneberg et al. – Supplement Fig. 7.**
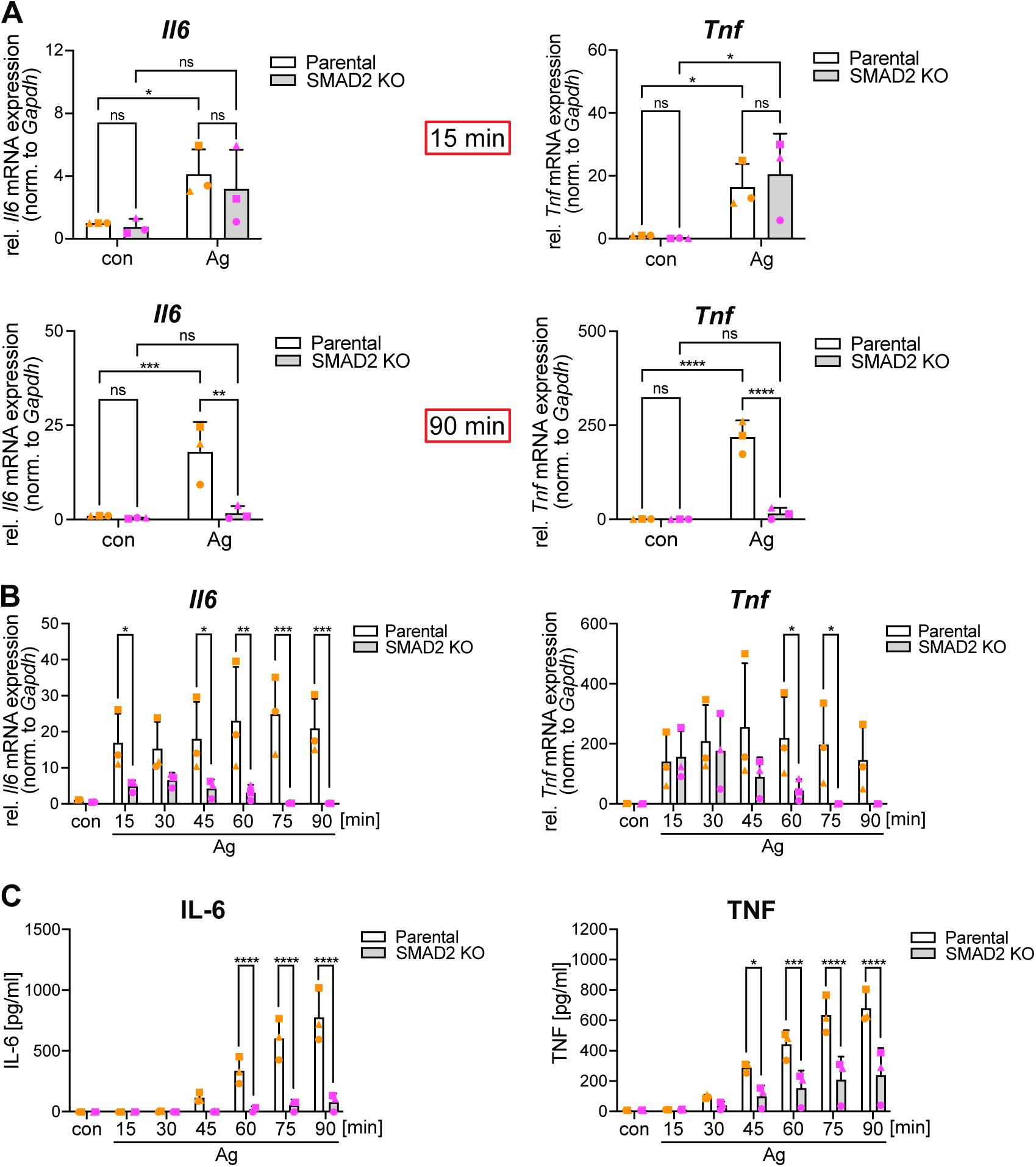
(A, B) The mRNA expression of *Il6* and *Tnf* was assessed by RT-qPCR in PMC-306 parental and S2KO cells in response to either 15 min and 90 min (A) or the indicated time points (B) of Ag. The mRNA expression was normalized to *Gapdh*. n=3 (C) The secretion of IL-6 and TNF from PMC-306 parental and S2KO cells in response to Ag for the indicated time points was measured by ELISA. n=3. Bar graphs are shown as mean + SD. *p<0.05, **p<0.01, ***p<0.001, ****p<0.0001 by two-way ANOVA and Fisher’s LSD test (A) or two-way ANOVA and Tukey’s multiple comparison test (B, C).

**Bronneberg et al. – Supplement Fig. 8.**
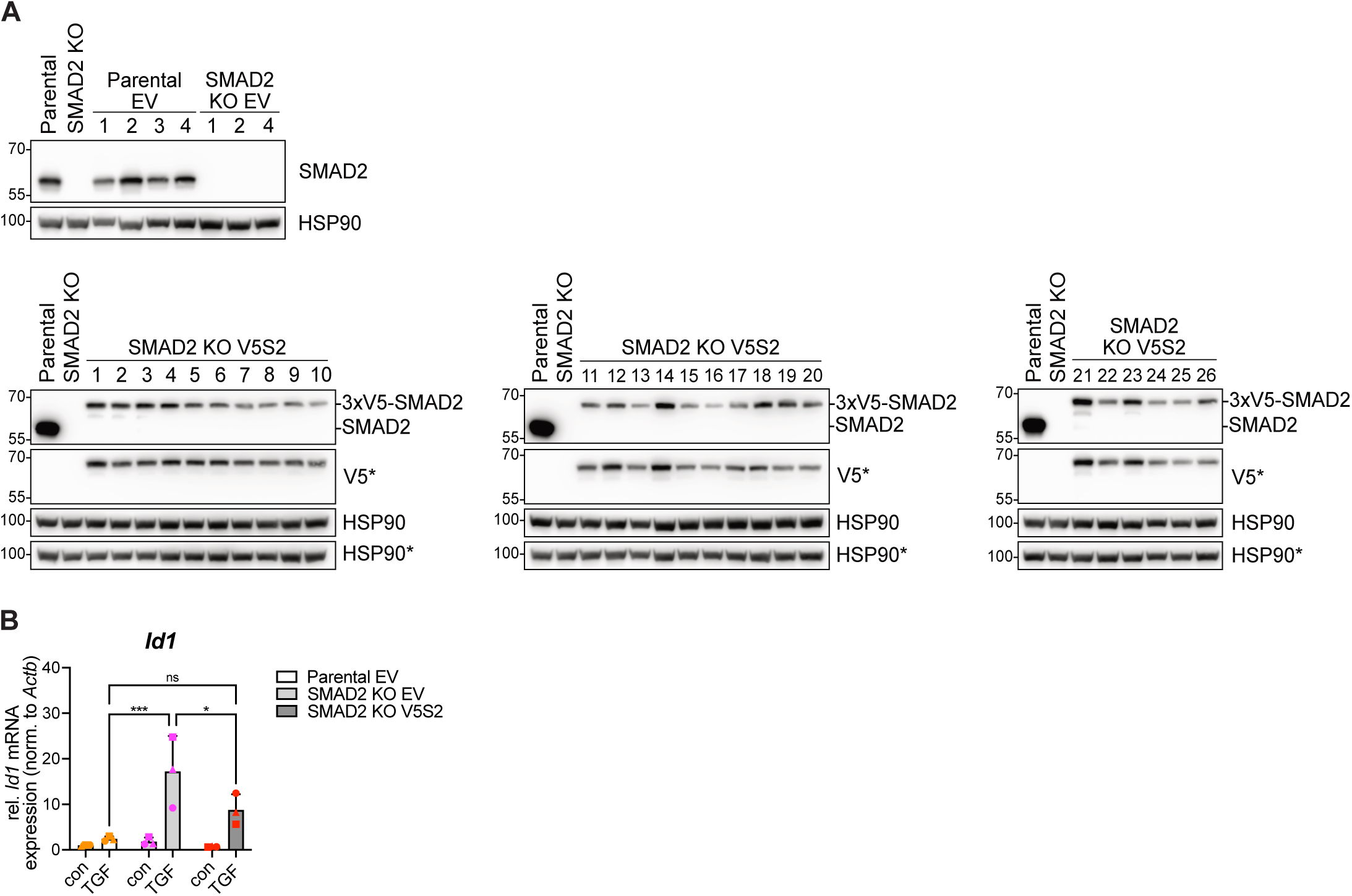
(A) Representative Western blots for SMAD2 and V5-tag in protein extracts from single-colony PMC-306 parental empty vector (EV), S2KO EV and S2KO 3xV5-Smad2 (V5S2) cells. HSP90 served as loading control. Asterisks indicate detection on the same membrane. (B) The mRNA expression of *Id1* was evaluated by RT-qPCR in PMC-306 parental EV, S2KO EV, and S2KO V5S2 cells after 90 min with or without TGF-β treatment. The mRNA expression was normalized to *Actb*. Three independent experiments are presented as mean + SD. *p<0.05, **p<0.01, ***p<0.001 by two-way ANOVA and Tukey’s multiple comparison test.

## References

1. St John AL, Rathore APS, Ginhoux F. New perspectives on the origins and heterogeneity of mast cells. Nat Rev Immunol 2023;23:55–68.

2. Valent P, Akin C, Hartmann K, Nilsson G, Reiter A, Hermine O et al. Mast cells as a unique hematopoietic lineage and cell system: From Paul Ehrlich’s visions to precision medicine concepts. Theranostics 2020;10:10743–10768.

3. Huber M, Cato ACB, Ainooson GK, Freichel M, Tsvilovskyy V, Jessberger R et al. Regulation of the pleiotropic effects of tissue-resident mast cells. J Allergy Clin Immunol 2019;144:S31–S45.

4. Galli SJ, Gaudenzio N, Tsai M. Mast Cells in Inflammation and Disease: Recent Progress and Ongoing Concerns. Annu Rev Immunol 2020;38:49–77.

5. Huber M. Activation/Inhibition of mast cells by supra-optimal antigen concentrations. Cell Commun Signal 2013;11:7.

6. Haque TT, Frischmeyer-Guerrerio PA. The Role of TGFbeta and Other Cytokines in Regulating Mast Cell Functions in Allergic Inflammation. Int J Mol Sci 2022;23. doi:10.3390/ijms231810864

7. Derakhshan T, Samuchiwal SK, Hallen N, Bankova LG, Boyce JA, Barrett NA et al. Lineage-specific regulation of inducible and constitutive mast cells in allergic airway inflammation. J Exp Med 2021;218. doi:10.1084/jem.20200321

8. Derakhshan T, Hollers E, Perniss A, Ryan T, McGill A, Hacker J et al. Human intraepithelial mast cell differentiation and effector function are directed by TGF-beta signaling. J Clin Invest 2025;135. doi:10.1172/JCI174981

9. Massague J, Sheppard D. TGF-beta signaling in health and disease. Cell 2023;186:4007–4037.

10. Heldin CH, Moustakas A. Signaling Receptors for TGF-beta Family Members. Cold Spring Harb Perspect Biol 2016;8. doi:10.1101/cshperspect.a022053

11. Derynck R, Budi EH. Specificity, versatility, and control of TGF-beta family signaling. Sci Signal 2019;12. doi:10.1126/scisignal.aav5183

12. Aashaq S, Batool A, Mir SA, Beigh MA, Andrabi KI, Shah ZA. TGF-beta signaling: A recap of SMAD-independent and SMAD-dependent pathways. J Cell Physiol 2022;237:59–85.

13. Nakao A, Afrakhte M, Moren A, Nakayama T, Christian JL, Heuchel R et al. Identification of Smad7, a TGFbeta-inducible antagonist of TGF-beta signalling. Nature 1997;389:631–635.

14. Kang Y, Chen CR, Massague J. A self-enabling TGFbeta response coupled to stress signaling: Smad engages stress response factor ATF3 for Id1 repression in epithelial cells. Mol Cell 2003;11:915–926.

15. Lebrin F, Goumans MJ, Jonker L, Carvalho RL, Valdimarsdottir G, Thorikay M et al. Endoglin promotes endothelial cell proliferation and TGF-beta/ALK1 signal transduction. EMBO J 2004;23:4018–4028.

16. Meurer SK, Bronneberg G, Penners C, Kauffmann M, Braunschweig T, Liedtke C et al. TGF-beta1 Induces Mucosal Mast Cell Genes and is Negatively Regulated by the IL-3/ERK1/2 Axis. Cell Commun Signal 2025;23:76.

17. Macias MJ, Martin-Malpartida P, Massague J. Structural determinants of Smad function in TGF-beta signaling. Trends Biochem Sci 2015;40:296–308.

18. Kretzschmar M, Doody J, Massague J. Opposing BMP and EGF signalling pathways converge on the TGF-beta family mediator Smad1. Nature 1997;389:618–622.

19. Liang J, Zhou Y, Zhang N, Wang D, Cheng X, Li K et al. The phosphorylation of the Smad2/3 linker region by nemo-like kinase regulates TGF-β signaling. Journal of Biological Chemistry 2021;296. doi:10.1016/j.jbc.2021.100512

20. Kamato D, Burch M, Zhou Y, Mohamed R, Stow JL, Osman N et al. Individual Smad2 linker region phosphorylation sites determine the expression of proteoglycan and glycosaminoglycan synthesizing genes. Cell Signal 2019;53:365–373.

21. Babaahmadi-Rezaei H, Kheirollah A, Rashidi M, Seif F, Niknam Z, Zamanpour M. EGF Receptor Transactivation by Endothelin-1 Increased CHSY-1 Mediated by NADPH Oxidase and Phosphorylation of ERK1/2. Cell J 2021;23:510–515.

22. Capellmann S, Sonntag R, Schuler H, Meurer SK, Gan L, Kauffmann M et al. Transformation of primary murine peritoneal mast cells by constitutive KIT activation is accompanied by loss of Cdkn2a/Arf expression. Front Immunol 2023;14:1154416.

23. Rostam MA, Shajimoon A, Kamato D, Mitra P, Piva TJ, Getachew R et al. Flavopiridol Inhibits TGF-beta-Stimulated Biglycan Synthesis by Blocking Linker Region Phosphorylation and Nuclear Translocation of Smad2. J Pharmacol Exp Ther 2018;365:156–164.

24. Pemberton AD, Brown JK, Wright SH, Knight PA, Miller HR. The proteome of mouse mucosal mast cell homologues: the role of transforming growth factor beta1. Proteomics 2006;6:623–631.

25. Zhao W, Gomez G, Yu SH, Ryan JJ, Schwartz LB. TGF-beta1 attenuates mediator release and de novo Kit expression by human skin mast cells through a Smad-dependent pathway. J Immunol 2008;181:7263–7272.

26. Deng Z, Fan T, Xiao C, Tian H, Zheng Y, Li C et al. TGF-beta signaling in health, disease, and therapeutics. Signal Transduct Target Ther 2024;9:61.

27. Yang J, Davies RJ, Southwood M, Long L, Yang X, Sobolewski A et al. Mutations in bone morphogenetic protein type II receptor cause dysregulation of Id gene expression in pulmonary artery smooth muscle cells: implications for familial pulmonary arterial hypertension. Circ Res 2008;102:1212–1221.

28. Ramachandran A, Vizan P, Das D, Chakravarty P, Vogt J, Rogers KW et al. TGF-beta uses a novel mode of receptor activation to phosphorylate SMAD1/5 and induce epithelial-to-mesenchymal transition. Elife 2018;7. doi:10.7554/eLife.31756

29. Yu PB, Deng DY, Lai CS, Hong CC, Cuny GD, Bouxsein ML et al. BMP type I receptor inhibition reduces heterotopic [corrected] ossification. Nat Med 2008;14:1363–1369.

30. Martin-Malpartida P, Batet M, Kaczmarska Z, Freier R, Gomes T, Aragon E et al. Structural basis for genome wide recognition of 5-bp GC motifs by SMAD transcription factors. Nat Commun 2017;8:2070.

31. Benchabane H, Wrana JL. GATA-and Smad1-dependent enhancers in the Smad7 gene differentially interpret bone morphogenetic protein concentrations. Mol Cell Biol 2003;23:6646–6661.

32. Goumans MJ, Valdimarsdottir G, Itoh S, Lebrin F, Larsson J, Mummery C et al. Activin receptor-like kinase (ALK)1 is an antagonistic mediator of lateral TGFbeta/ALK5 signaling. Mol Cell 2003;12:817–828.

33. Sanchez-Duffhues G, Williams E, Goumans MJ, Heldin CH, Ten Dijke P. Bone morphogenetic protein receptors: Structure, function and targeting by selective small molecule kinase inhibitors. Bone 2020;138:115472.

34. Daly AC, Randall RA, Hill CS. Transforming growth factor beta-induced Smad1/5 phosphorylation in epithelial cells is mediated by novel receptor complexes and is essential for anchorage-independent growth. Mol Cell Biol 2008;28:6889–6902.

35. Liu IM, Schilling SH, Knouse KA, Choy L, Derynck R, Wang XF. TGFbeta-stimulated Smad1/5 phosphorylation requires the ALK5 L45 loop and mediates the pro-migratory TGFbeta switch. EMBO J 2009;28:88–98.

36. Chen HB, Shen J, Ip YT, Xu L. Identification of phosphatases for Smad in the BMP/DPP pathway. Genes Dev 2006;20:648–653.

37. Schmierer B, Tournier AL, Bates PA, Hill CS. Mathematical modeling identifies Smad nucleocytoplasmic shuttling as a dynamic signal-interpreting system. Proc Natl Acad Sci U S A 2008;105:6608–6613.

38. Duan X, Liang YY, Feng XH, Lin X. Protein serine/threonine phosphatase PPM1A dephosphorylates Smad1 in the bone morphogenetic protein signaling pathway. J Biol Chem 2006;281:36526–36532.

39. Knockaert M, Sapkota G, Alarcon C, Massague J, Brivanlou AH. Unique players in the BMP pathway: small C-terminal domain phosphatases dephosphorylate Smad1 to attenuate BMP signaling. Proc Natl Acad Sci U S A 2006;103:11940–11945.

40. Wrighton KH, Lin X, Feng XH. Phospho-control of TGF-beta superfamily signaling. Cell Res 2009;19:8–20.

41. Kamato D, Little PJ. Smad2 linker region phosphorylation is an autonomous cell signalling pathway: Implications for multiple disease pathologies. Biomed Pharmacother 2020;124:109854.

42. Kretzschmar M, Doody J, Timokhina I, Massague J. A mechanism of repression of TGFbeta/ Smad signaling by oncogenic Ras. Genes Dev 1999;13:804–816.

43. Afroz R, Zhou Y, Little PJ, Xu S, Mohamed R, Stow J et al. Toll-like Receptor 4 Stimulates Gene Expression via Smad2 Linker Region Phosphorylation in Vascular Smooth Muscle Cells. ACS Pharmacol Transl Sci 2020;3:524–534.

44. Rostam MA, Kamato D, Piva TJ, Zheng W, Little PJ, Osman N. The role of specific Smad linker region phosphorylation in TGF-beta mediated expression of glycosaminoglycan synthesizing enzymes in vascular smooth muscle. Cell Signal 2016;28:956–966.

45. Ferrell Jr JE. Self-perpetuating states in signal transduction: Positive feedback, double-negative feedback and bistability. Curr Opin Chem Biol 2002;6:140–148.

46. Meurer SK, Ness M, Weiskirchen S, Kim P, Tag CG, Kauffmann M et al. Isolation of Mature (Peritoneum-Derived) Mast Cells and Immature (Bone Marrow-Derived) Mast Cell Precursors from Mice. PLoS One 2016;11:e0158104.

## References

1. Meurer SK, Ness M, Weiskirchen S, Kim P, Tag CG, Kauffmann M et al. Isolation of Mature (Peritoneum-Derived) Mast Cells and Immature (Bone Marrow-Derived) Mast Cell Precursors from Mice. PLoS One 2016;11:e0158104.

2. Karasuyama H, Melchers F. Establishment of mouse cell lines which constitutively secrete large quantities of interleukin 2, 3, 4 or 5, using modified cDNA expression vectors. Eur J Immunol 1988;18:97–104.

3. Conant D, Hsiau T, Rossi N, Oki J, Maures T, Waite K et al. Inference of CRISPR Edits from Sanger Trace Data. CRISPR J 2022;5:123–130.

4. Ewels PA, Peltzer A, Fillinger S, Patel H, Alneberg J, Wilm A et al. The nf-core framework for community-curated bioinformatics pipelines. Nat Biotechnol 2020;38:276–278.

5. Di Tommaso P, Chatzou M, Floden EW, Barja PP, Palumbo E, Notredame C. Nextflow enables reproducible computational workflows. Nat Biotechnol 2017;35:316–319.

6. Krueger F, James F, Ewels P, Afyounian E, Schuster B. FelixKrueger/TrimGalore: v0.6.7 - DOI via Zenodo. 2021. 10.5281/zenodo.5127899

7. Dobin A, Davis CA, Schlesinger F, Drenkow J, Zaleski C, Jha S et al. STAR: ultrafast universal RNA-seq aligner. Bioinformatics 2013;29:15–21.

8. Patro R, Duggal G, Love MI, Irizarry RA, Kingsford C. Salmon provides fast and bias-aware quantification of transcript expression. Nat Methods 2017;14:417–419.

9. Love MI, Huber W, Anders S. Moderated estimation of fold change and dispersion for RNA-seq data with DESeq2. Genome Biol 2014;15:550.

10. Meurer SK, Bronneberg G, Penners C, Kauffmann M, Braunschweig T, Liedtke C et al. TGF-beta1 Induces Mucosal Mast Cell Genes and is Negatively Regulated by the IL-3/ERK1/2 Axis. Cell Commun Signal 2025;23:76.

11. Pfaffl MW. A new mathematical model for relative quantification in real-time RT-PCR. Nucleic Acids Res 2001;29:e45.

